# Metabolic Switch in Endocrine Resistant Estrogen Receptor Positive Breast Cancer

**DOI:** 10.1101/2024.12.28.630631

**Authors:** Heather M. Brechbuhl, Amy Han, Kiran Vinod Paul, Travis Nemkov, Srinivas Ramachandran, Ashley Ward, Britta M. Jacobsen, Kirk Hansen, Carol A. Sartorius, Angelo D’Alessandro, Peter Kabos

## Abstract

**Purpose:** The development of endocrine resistance remains a significant challenge in the clinical management of estrogen receptor-positive (**ER+**) breast cancer. Metabolic reprogramming is a prominent component of endocrine resistance and a potential therapeutic intervention point. However, a limited understanding of which metabolic changes are conserved across the heterogeneous landscape of ER+ breast cancer or how metabolic changes factor into ER DNA binding patterns hinder our ability to target metabolic adaptation as a treatment strategy. This study uses dimethyl fumarate (**DMF**) to restore tamoxifen (**Tam**) and fulvestrant (**Fulv**) sensitivity in endocrine-resistant cell lines and investigates how metabolic changes influence ER DNA-binding patterns.

**Experimental Design:** To address the challenge of metabolic adaptation in anti-endocrine resistance, we generated Tam and Fulv resistance in six ER+ breast cancer (**BC**) cell lines, representing ductal (MCF7, T47D, ZR75-1, and UCD12), lobular (MDA-MB-134--VI), and HER2 amplified (BT474) BC molecular phenotypes. Metabolomic profiling, RNA sequencing, proteomics, and CUT&RUN assays were completed to characterize metabolic shifts, transcriptional and protein changes, and ER DNA-binding patterns in resistant cells. Dimethyl fumarate was assessed for its ability to reverse Tam and Fulv resistance, restore tricarboxylic acid cycle (**TCA**) cycle function, and restore parental cell (endocrine sensitive) ER DNA binding patterns.

**Results:** Tamoxifen-resistant (TamR) and fulvestrant-resistant (FulvR) cells exhibited disrupted TCA cycle activity, reduced glutathione levels, and altered nucleotide and amino acid metabolism. DMF treatment replenished TCA cycle intermediates and reversed resistance in both TamR and FulvR cells. DMF also increased mevalonate pathway enzyme expression in both TamR and FulvR cells, with TamR cells upregulating enzymes in the cholesterol synthesis phase and FulvR enhancing enzymes in the early part of the pathway. DMF restored ER DNA-binding patterns in TamR cells to resemble parental cells, re-sensitizing them to Tam. In FulvR cells, DMF reversed resistance by modulating ER-cofactor interactions but did not restore parental ER DNA-binding signatures.

**Conclusions:** Our findings provide new insights into how metabolic reprogramming affects ER DNA-binding activity in endocrine-resistant breast cancer. We demonstrate how altering metabolism can reprogram ER signaling and influence resistance mechanisms by targeting metabolic vulnerabilities, such as TCA cycle disruptions. Additionally, our data provide a comprehensive metabolomic, RNA-seq, and CUT&RUN data set relevant to tumor metabolic adaptation leading to acquired endocrine resistance in highly utilized ER+ breast cancer cell lines. This study improves our understanding of how metabolic states alter ER function in endocrine-resistant breast cancer.

## INTRODUCTION

Metabolic dysfunction is a hallmark found in progressing breast cancer (**BC**) and contributes to acquired drug resistance by reducing the effectiveness of anti-endocrine therapies. Such resistance is a significant factor in disease progression and leads to unfavorable patient outcomes. Targeting tumor cell metabolism is a key area of new drug development. Several FDA-approved drugs for breast cancer impact metabolism, such as everolimus, which targets the mTOR pathway, and alpelisib and taselisib, which target PI3K signaling. Even common chemotherapies such as 5-fluorouracil, gemcitabine, and methotrexate affect metabolism by interfering with nucleotide synthesis. Even with the success of these drugs, we still lack an understanding of how metabolic adaptation alters the effect of anti-endocrine therapeutics, which represent the standard treatment for estrogen receptor-positive (**ER+**) BC.

The estrogen receptor has a vital role in regulating the metabolism of ER+ BC by promoting the uptake and utilization of nutrients essential for breast cancer establishment, growth, and spread. Anti-endocrine therapy is aimed at blocking the ability of the estrogen receptor to bind DNA and thereby inhibit its pro-tumor effects. Three approaches have been used to inhibit estrogen receptor activity: aromatase inhibitors (i.e., exemestane) block estrogen production; selective estrogen receptor modulators (i.e., tamoxifen) compete with estrogen for binding of the estrogen receptor active site, and selective estrogen receptor degraders (i.e., fulvestrant) mark the estrogen receptor for degradation. Acquired resistance to anti-endocrine therapies is frequent, and metabolic adaptation is known to promote acquired resistance. However, little is known about how metabolic dysfunction affects estrogen receptor activity.

In this study, we addressed the challenge of metabolic adaptation in anti-endocrine resistance by generating tamoxifen (**Tam**) and fulvestrant (**Fulv**) resistance in six ER+ breast cancer (**BC**) cell lines, representing ductal (MCF7, T47D, ZR75-1, and UCD12), lobular (MDA-MB-134-VI), and HER2 amplified (BT474) molecular phenotypes. Using an-omics approach, we determined which metabolic pathways were conserved within and between Tam-resistant (**TamR**) and Fulv-resistant (**FulvR**) cell lines. We found that both TamR and FulvR BC cells had a disruption in the TCA cycle that could be resolved by treating the cells with the clinically approved drug dimethyl fumarate (**DMF**). Finally, we characterized the ability of estrogen receptor binding to DNA in DMF-treated TamR and FulvR cells. We found that compared to BC cells sensitive to Tam and Fulv treatment (parental cells), TamR and FulvR cells had significantly different estrogen receptor DNA binding signatures. Treating TamR cells with DMF restored large portions of the estrogen receptor DNA binding pattern back to the parental pattern and re-sensitized the TamR cells to Tam treatment. DMF treatment also restored the sensitivity of FulvR cells to Fulv treatment. Still, this re-sensitization was not associated with the significant restoration of parental patterns for estrogen receptor DNA binding. Instead, the restoration activated the mevalonate pathway and was associated with interactions with transcription factors that are complex with ER to exert their effects.

## MATERIALS and METHODS

### Development of BC cell line resistance

ER+ breast cancer cell lines (BT474, MCF-7, MDA-MB-134-VI, T47D, ZR75-1, and UCD12) were exposed to escalating doses of Tam or Fulv over 12 months to establish endocrine resistance. The starting dose for Fulv and Tam were 10 times lower than the effective dose, (the dose capable of significantly inhibiting growth)for each compound, 1nM and 10nM, respectively. Both compounds were dissolved in 100% DMSO to make stock concentrations that resulted in the final cell culture dose being less than 10% DMSO. All lines were established to 1 µM resistance for Tam and Fulv, except for T47D and UCD12 cells, which were established to 60 nM resistance for Fulv. Cell lines were assessed by metabolomics and bulk RNA sequencing to compare metabolic changes associated with the development of Tam and Fulv resistance. All cell lines tested mycoplasma negative before use in any downstream application using the Universal Mycoplasma PCR-Based detection kit from ATCC.

### Sample Processing for Metabolomics, RNA sequencing, and Proteomics

FulvR and TamR cell lines and their parental (drug-sensitive) equivalent cells were split and allowed to grow for 72 hours, reaching approximately 70% confluence in the presence of the respective drug or DMSO control. The media was replaced with drug-free media overnight to wash out all remnants of compounds that could interfere with downstream assays. Cells were trypsinized and pelleted by centrifugation, washed 2 times in ice-cold phosphate-buffered saline, counted by hemocytometer, and adjusted to make pellets of at least 1 million cells per pellet. Next, cells were pelleted by centrifugation and snap-frozen on dry ice for storage at −80°C until ready for downstream metabolomic analysis. Metabolomic analysis was done through the University of Colorado Mass Spectrometry Metabolomics Shared Resource Facility. RNA sequencing libraries were made using the TruSeq® Stranded mRNA Library Prep Kit, and the University of Colorado Cancer Center Genomics Shared Resource sequenced the libraries. Proteomics analysis was completed through the University of Colorado Mass Spectrometry Proteomics Shared Resource Facility.

### DMF Treatment and Live Cell Imaging

For live cell imaging assays, cells were plated in a 96-well plate at approximately 25 percent confluence in standard media and treated with either 10nM Fulv, 100nM Tam, 50µM DMF, vehicle control (10% DMSO), or a combination of Tam + DMF or Fulv + DMF. Images were automatically taken every 4 hours over 72 hours in a BioTek BioSpa Automated Incubator (Agilent, Santa Clara, CA). Cell proliferation was quantified using Aligent’s Gen5 image analysis tools.

### ER Cut&Run and Motif Analysis

After 72 hours of treatment with EtOH or 50 µM DMF, cells were washed twice with phosphate-buffered saline and harvested using a cell lifter. Each sample contained 5 x 10^5^ cells for downstream processing according to the protocol adapted from Skene P et al. [1] and detailed in Han A. et al. [2]. The Ovation Ultralow System V2 kit by Tecan was used to generate CUT&RUN libraries according to the kit instructions and amplification conditions: 98°C for 45s, (98°C for 15s, 60°C for 10s) x 13 cycles, 72°C for 1 min, followed by 4°C hold. A detailed protocol is found in the kit and is described by Han A. et al. [2]. Libraries were submitted to the Genomics and Microarray Shared Resource at the University of Colorado Anschutz, and 10 million paired-end reads per sample were detected on a NovaSEQ6000 (Illumina, San Diego, CA). Data were processed and analyzed according to the detailed description provided by Han A. et al. for defining the CUT&RUN score, differential peak calling, peak clustering, motif discovery, and fractions [2]. Motif analysis was performed using HOMER and FIMO.

### Joint Pathway Analysis

We determined the log2 fold-change between the parent and TamR or FulvR equivalent cell line for each metabolite and coding transcript. Because the metabolic profiling was limited to 193 identified metabolites in every cell line, but RNA sequencing identified several thousand coding genes, we did not want the RNA sequencing to completely wash out the metabolite results from the Joint Pathway Analysis (JPA) tool in MetaboAnalyst. Therefore, we included all metabolites with log2 fold-change values as input to JPA. However, we narrowed our coding gene list from RNAseq by inputting transcripts with −0.6 > log2 fold-change > 0.6 and a false discovery rate < 0.05. Reported pathways had a p-value < 0.019, a false discovery rate < 0.055, and a composite score > 1.0. The impact score output from MetaboAnalyst is weighted based on the centrality of the metabolite in a pathway, with metabolites that have a more substantial influence on pathway activity being given more weight than those metabolites in the pathway less central to the activity of the given pathway. To calculate the composite score, we used the calculation −Log10(FDR)*(impact score)^2^.

### Statistics

MetaboAnalyst was used to normalize, range scale, and complete partial-least-squares discriminant analysis. GraphPad Prism was version 9.5.1 analytical software used for determining statistical significance throughout the paper. We used unpaired two-tailed t-tests with assumptions of parametric Gaussian distribution and equal standard deviations for single comparisons. We used ordinary one-way ANOVA analysis with Tukey multiple comparisons tests for multiple comparisons. K-means clustering of the CUT&RUN data was completed according to the method described by Lloyd, S. [3]. All experiments were conducted using at least triplicate replicates and the biological replicates represented by each cell line. Outliers were identified using GraphPad Prism Outlier ROUT function with Q = 1%. All data are presented as mean +/− the standard deviation.

## RESULTS

### Metabolic and transcriptomic changes define breast cancer drug resistance

To address the challenge of metabolic adaptation in anti-endocrine resistance, we used a dose escalation protocol (see Methods) to generate a final set of ER+ **BC** cell lines resistant to Tam or Fulv. The cells were considered resistant to the drug once they survived in a dose 10 times higher than the effective dose in a sensitive ER+ BC cell line (**Figure 1A-B**). Matching drug-sensitive (**parental**) cell lines were maintained alongside the resistant cells and demonstrated decreased growth in response to 100 nM Tam or 10 nM Fulv treatment.

**Figure 1.**
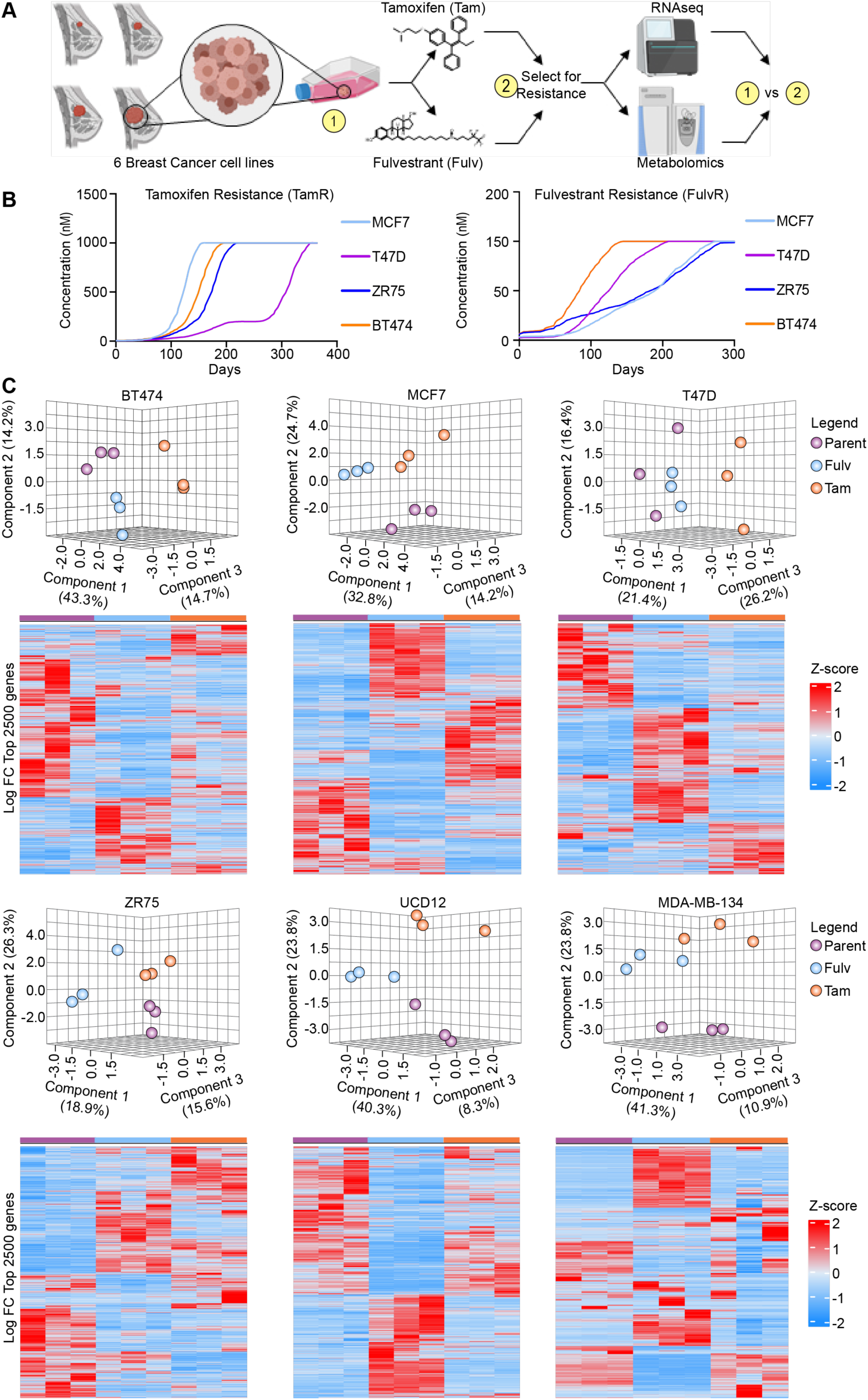
Metabolomic characteristics of endocrine-resistant cells. Resistance to Tam or Fulv results in a clear distinction between cell types based on their endocrine responsiveness. (**A**) Workflow diagram for generation and analysis of tamoxifen (Tam) and fulvestrant (Fulv) resistant cell lines. ER+ cell lines were grown over a 12-month period and exposed to escalating doses of (**B**) TAM or Fulv. (**C**) Partial Least Squares Discriminant Analysis (PLS-DA) analysis of the metabolomic profile for individual cell lines and treatment groups (parent, TAM-resistant, and Fulv-resistant) paired with the heat map of the top 2500 protein-coding transcripts from RNA-seq in each cell line and treatment group.

We analyzed the metabolomes and transcriptomes of the resistant cell lines to determine how TamR and FulvR cells were distinguished from the parental cells. Using MetaboAnalyst, we normalized the metabolomes of FulvR and TamR cell lines to their respective parental line and generated a partial-least squares discriminant analysis (**PLS-DA**) for each cell type. PLS-DA demonstrated that within each cell type, FulvR and TamR were easily discerned from the parental cells (**Figure 1C**). Additionally, hierarchal cluster analysis (HCA) generated from the top 2500 transcripts for coding genes in each cell type demonstrated that the transcriptomes of FulvR, TamR, and parental cells are distinct from each other (**Figure 1C**).

### Endocrine-resistant BCCs show altered building blocks and disrupted redox balance

To determine shared metabolic traits associated with TamR and FulvR, we analyzed BC cells as a function of treatment resistance compared to parental lines. This analysis revealed significant changes in nucleotide metabolism (**Figure 2**). Specifically, both resistance mechanisms exhibited shared increases in purine abundance, with elevated levels of guanine, guanosine, and adenosine. These trends may be fueled by upregulated nucleotide salvage pathways, as evidenced by shared elevations in inosine and hypoxanthine, which are intermediates of purine salvage.

**Figure 2.**
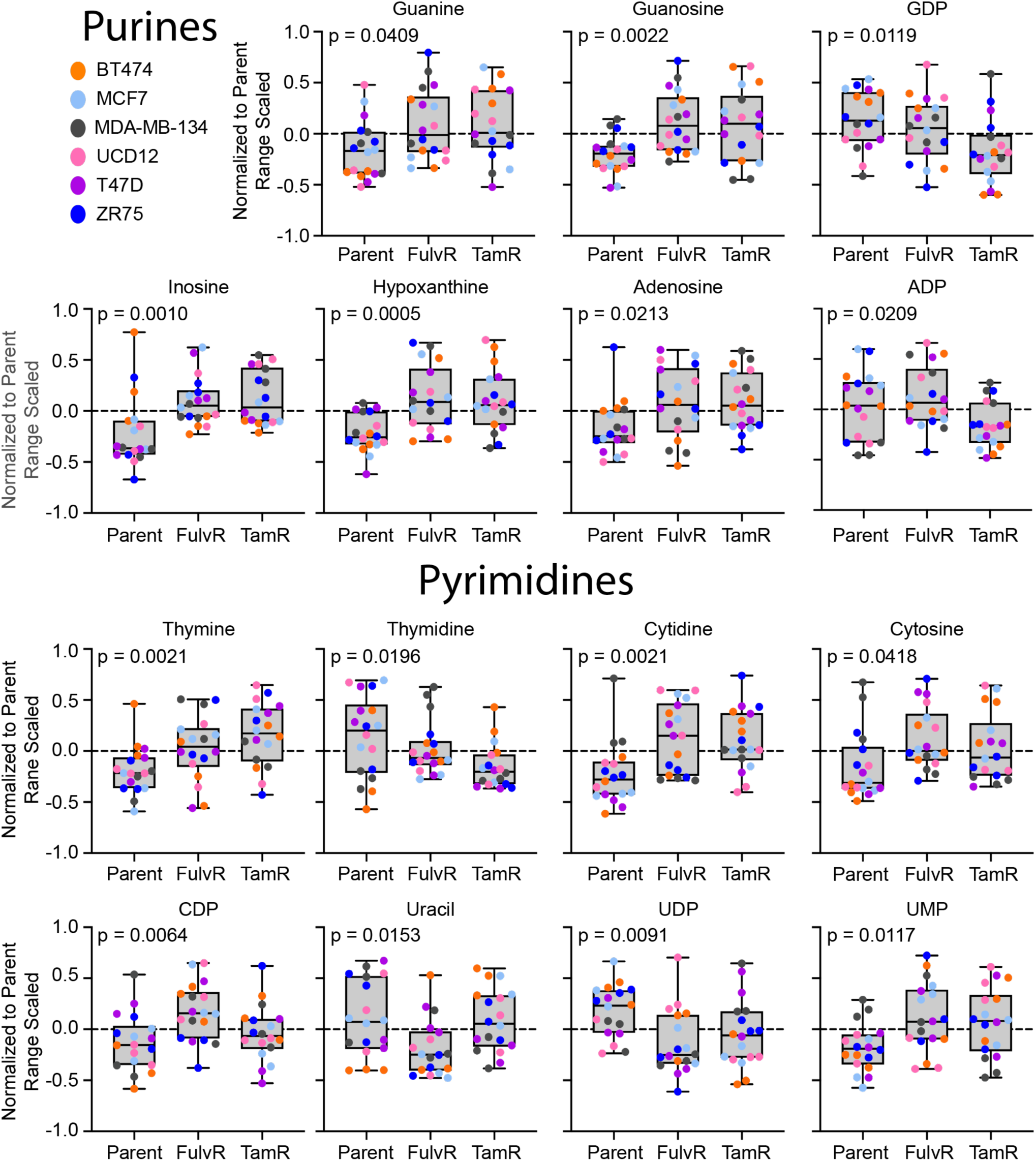
Endocrine-resistant ER+ breast cancer cell lines have alterations in nucleotide metabolism. Box and whiskers plots of significantly changed purine and pyrimidine levels in TamR and FulvR cells compared to parental. Statistical analysis is 1-way ANOVA, and significance is p < 0.05.

Interestingly, while guanosine diphosphate (**GDP**) and adenosine diphosphate (**ADP**) levels remained unchanged in FulvR cells compared to parental lines, they were notably decreased in TamR cells. Despite these changes in GDP and ADP, adenosine triphosphate (**ATP**) and adenosine monophosphate (**AMP**) levels did not significantly differ between any of the groups. The reduction in GDP and ADP, specifically in TamR cells, combined with stable ATP levels, suggests impairment in the adenine nucleotide pool turnover, which could affect the overall energy balance. These alterations point to a unique metabolic vulnerability in TamR cells related to nucleotide handling and energy change, potentially impacting their ability to maintain proper energy homeostasis during the acquisition of Tam resistance.

Shared responses in pyrimidine metabolism were observed between TamR and FulvR cells, with both groups showing increased levels of cytidine and cytosine compared to parental cells. Both resistance types also displayed elevated thymine levels, although thymidine was lower than parental lines, indicating potential alterations in pyrimidine-nucleoside phosphorylation. This trend was particularly pronounced in FulvR cells, which also showed higher levels of cytidine diphosphate (**CDP**) compared to both parental and TamR cells. Interestingly, FulvR cells maintained significantly lower levels of uracil, while both TamR and FulvR cells exhibited decreased uridine diphosphate (**UDP**) and increased uridine monophosphate (**UMP**). These findings suggest distinct disruptions in pyrimidine metabolism, particularly in the phosphorylation states of uridine nucleotides, which may play a role in the metabolic adaptation associated with endocrine resistance.

Our results also revealed alterations in amino acid metabolism in both TamR and FulvR cells (**Figure 3**). In TamR cells, glycine and valine were significantly elevated, while alanine was significantly lower compared to parental cells. Arginine levels showed no significant difference, but histidine and lysine trended higher. In contrast, FulvR cells exhibited a slight trend towards increased glycine and valine levels, while alanine was significantly decreased. Arginine was significantly lower, and histidine trended lower with no significant changes in lysine levels. These alterations suggest differences in amino acid utilization between the two types of resistant cells, with specific modifications affecting pathways such as “Arginine biosynthesis,” “Alanine, aspartate and glutamate metabolism,” and “Glycine, serine and threonine metabolism” (**SFigure 1-6**) as identified by Joint Pathway Analysis using MetaboAnalyst.

**Figure 3.**
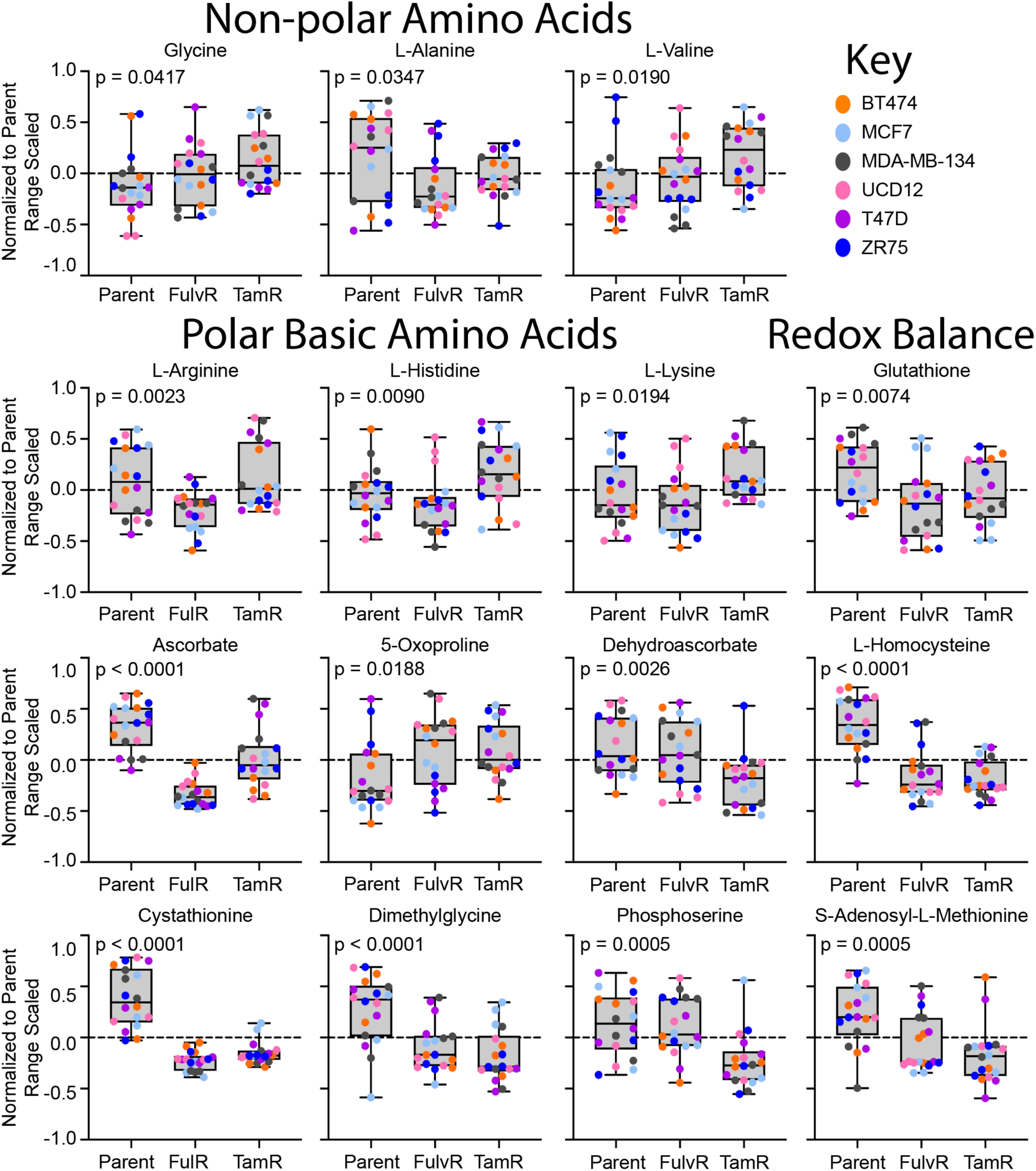
Endocrine-resistant ER+ breast cancer cell lines have several alterations in amino acid metabolism. Box and whiskers plots of significantly changed amino acid levels in TamR and FulvR cells compared to parental. Both cell lines demonstrated decreased levels of glutathione and intermediates of glutathione. Statistical analysis is 1-way ANOVA, and significance is p < 0.05.

Regarding redox balance, glutathione (**GSH**), a key cellular antioxidant, was significantly lower in both TamR and FulvR cells (**Figure 3**), indicating a compromised redox regulation. In TamR cells significant decreases in ascorbate and dehydroascorbate were observed, while FulvR cells had significantly reduced ascorbate with no change in dehydroascorbate. Both cell types exhibited significantly lower levels of L-homocysteine and cystathionine, two precursors for cysteine and GSH synthesis, further indicating disruptions in the methionine cycle. Interestingly, S-adenosyl-L-methionine (**SAM**), a key intermediate in the methionine cycle, was significantly lower in TamR cells and trended lower in FulvR cells, reinforcing the conclusion that methionine cycle activity is reduced. Additionally, dimethylglycine, another molecule involved in redox balance, was significantly decreased in both resistant cell types. Phosphoserine, which contributes to GSH peroxidase function, was significantly reduced in TamR cells but showed no significant changes in FulvR cells. Both cell types had elevated levels of 5-oxoproline, which indicates defects in cysteine synthesis from homocysteine. Finally, JPA also identified “Glutathione metabolism” as a significantly impacted pathway for all resistant cell lines, except for ZR75 FulvR cells, further supporting the conclusion that redox balance is disrupted in the resistant cells.

Taken together, these data suggest that the redox balance in TamR and FulvR cells is impaired due to disruptions in GSH metabolism, the methionine cycle, and cysteine availability. Given the connection between redox imbalance and energy production, we analyzed metabolic signatures related to glycolysis and the tricarboxylic acid (**TCA**) cycle to explore further the underlying metabolic vulnerabilities in TamR and FulvR cells.

### Metabolic shunting of TCA cycle intermediates in TamR and FulvR cells

TamR and FulvR cells exhibit significant alterations in the TCA cycle, suggesting a metabolic reprogramming that diverts resources away from efficient energy production. JPA identified enrichment of “Pyruvate metabolism” in TamR cells compared to FulvR and parental cells, which was supported in the metabolomic data, showing increased pyruvate levels (**SFigure 1-6** and **Figure 4**). Elevated pyruvate concentrations in the TamR cells indicate pyruvate accumulation upstream of the TCA cycle. This suggests a metabolic bottleneck or diversion of pyruvate away from its traditional entry point into mitochondrial respiration. Despite JPA enrichment for “Pyruvate metabolism” and elevated pyruvate levels in TamR cells compared to FulvR and parent, citrate levels were significantly decreased in TamR cells, suggesting that pyruvate was not being efficiently converted to citrate, again indicating a potential block of pyruvate shuttling into the TCA cycle. Furthermore, JPA also identified enrichment for the “Citrate cycle” in both TamR and FulvR cells (**SFigure 1-6**), indicating broader disruptions within mitochondrial metabolism. Compared to parental cells, FulvR cells had significantly elevated succinate levels, and TamR cells trended toward increased levels.

**Figure 4.**
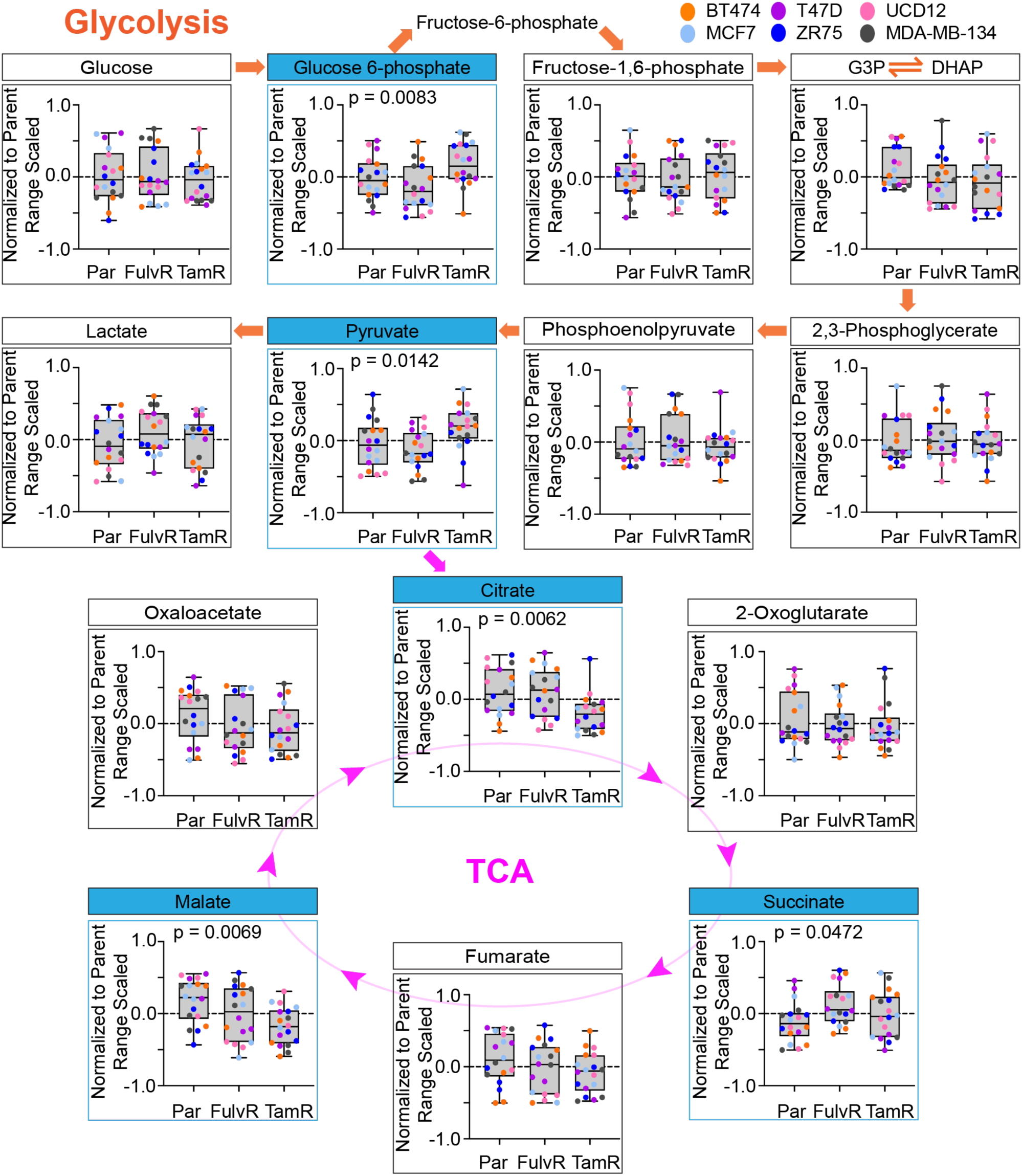
Endocrine-resistant ER+ breast cancer cells have altered glycolysis and decreased citrate cycle activity. Box and whiskers plots of metabolites in the glycolysis and TCA cycle. Statistical analysis is 1-way ANOVA, and significance is p < 0.05.

Downstream of succinate, fumarate levels trended lower in both TamR and FulvR compared to parent cells, suggesting a potential block or reduced activity of succinate dehydrogenase (complex II), which catalyzes the conversion of succinate to fumarate. Furthermore, malate and oxaloacetate levels also trended downwards in TamR and FulvR cells, with malate levels reaching significance in FulvR cells (**Figure 4**), suggesting a reduced flow through the latter parts of the TCA cycle and potential impairment of pathways that replenish TCA cycle intermediates. These findings indicate that both TamR and FulvR cells may have a blockade in mitochondrial complex II, resulting in decreased or rerouting of intermediates of the TCA cycle. We next aimed to reverse Tam and Fulv resistance by overcoming the apparent TCA blockade.

### TamR and FulvR cells are sensitive to dimethyl fumarate treatment

The compound Dimethyl Fumarate (**DMF**) can be metabolized into fumaric acid and has been shown to increase downstream metabolites of the TCA cycle [4]. We hypothesized that treating cells with DMF would bypass the blocked conversion of succinate to fumarate and force TCA progression through complex II. As expected, our results showed that TamR cells (including MCF7-, ZR75-, and UCD12-TamR) were resistant to 100nM Tam treatment alone by demonstrating significant growth despite the presence of the Tam. However, treatment with 50 µM DMF alone inhibited growth across all tested TamR lines (**Figure 5A and SFigure 7**), and combination treatment using 50 µM DMF and 100 nM Tam resulted in complete growth arrest of TamR cells.

**Figure 5.**
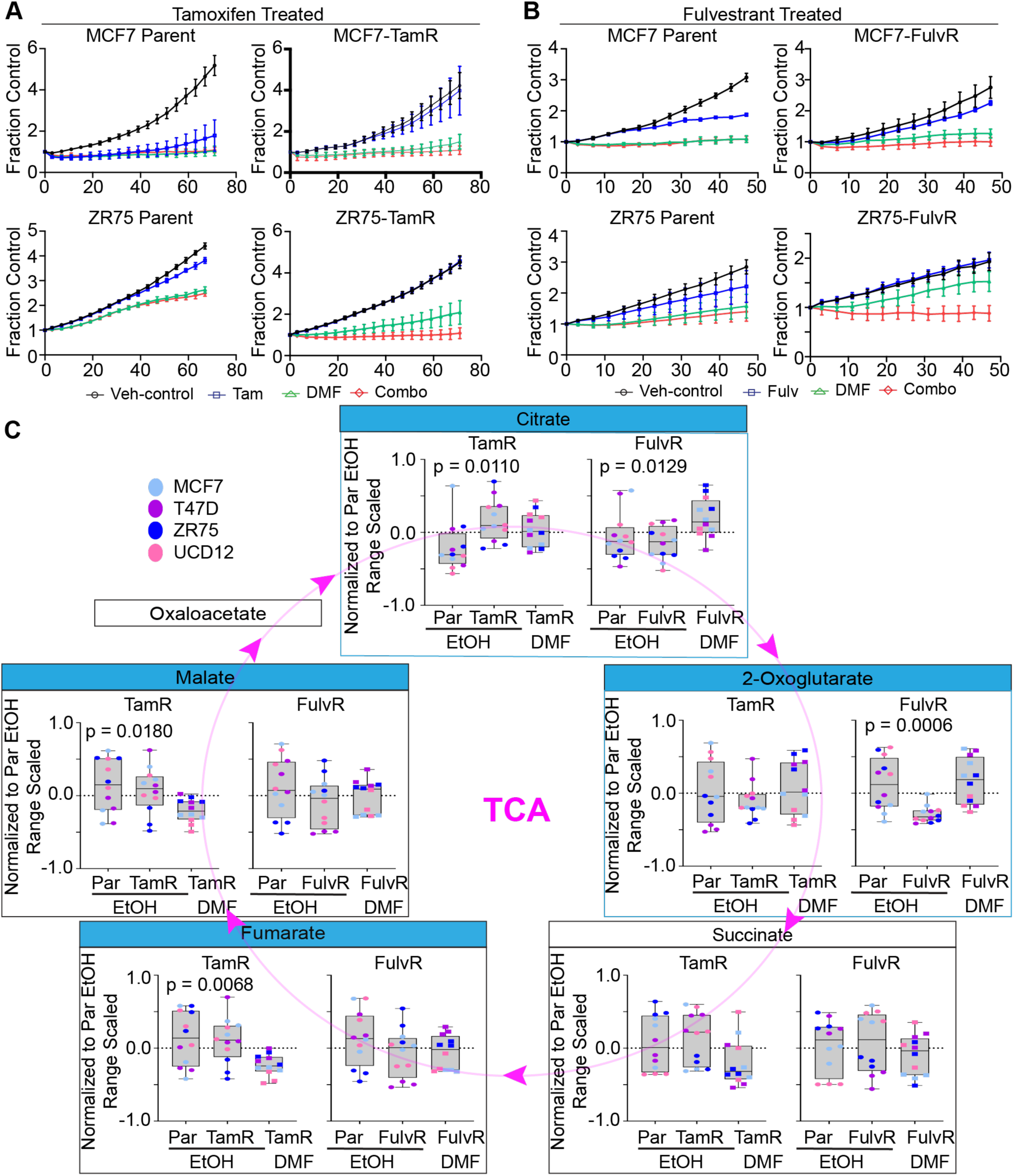
DMF reverses endocrine resistance by forcing cells through the TCA cycle. **(A)** Incucyte live cell growth curves show dimethyl fumarate (DMF) caused decreased cell growth alone and could reverse Tam and Fulv resistance when combined with DMF. **(B)** Box and whiskers plots of metabolites in the TCA cycle. Statistical analysis is 1-way ANOVA, and significance is p < 0.05.

Our findings also showed that FulvR cells (including MCF7- and ZR75-FulvR) proliferated well in the presence of 10 nM Fulv but had significant growth reduction when treated with 50 µM DMF alone, an effect strongest in FulvR-MCF7 cells (**Figure 5B**). Combination treatment with DMF and Fulv resulted in further growth reduction and was significantly better in FulvR-ZR75 cells than in DMF alone. DMF treatment also resulted in a significant increase in citrate and 2-Oxoglutarate, specifically in FulvR cells (**Figure 5C**), suggesting that DMF is more effective in restoring TCA cycle function in FulvR cells compared to TamR cells. This difference could be indicative of distinct metabolic dependencies between the resistance phenotypes and indicate that DMF may engage unique metabolic vulnerabilities between the two resistance types. Since DMF was sufficient to reverse endocrine resistance in both TamR and FulvR cells, we investigated whether the effects of DMF treatment could be attributed to changes in estrogen receptor (**ER**) activity.

### Following DMF treatment, the Mevalonate Pathway enzymes were enhanced in endocrine-resistant cells

The mevalonate pathway is crucial for cholesterol synthesis, a precursor for hormone steroids, including estradiol. Previous studies have reported alterations in lipid metabolism and increased HDL cholesterol levels associated with DMF treatment [5, 6]. Moreover, reduced expression or inhibition of 3-hydroxy-3-methylglutharyl-coenzyme A reductase (**HMG-CoAR**), the rate-limiting enzyme in the mevalonate pathway, has been correlated with reduced risk for ER+ breast cancer recurrence and reduced mortality [7–9].

We used our proteomic data to examine expression changes in the mevalonate pathway after DMF treatment in TamR and FulvR cells. The analysis detected 12 of the 13 enzymes in the mevalonate pathway, with 4 demonstrating increased expression in FulvR and 5 in TamR cells following DMF treatment (**SFigure 8-9**). Notably, DMF treatment in FulvR cells caused increased protein expression of the rate-limiting enzyme HMG-CoAR and enzymes in the early half of the pathway but not in the final enzymes responsible for cholesterol synthesis (**SFigure 8**). However, TamR cells upregulated enzyme expression in the last half of the pathway responsible for cholesterol synthesis but did not upregulate the rate-limiting enzyme HMG-CoAR (**SFigure 9**). Previous research has demonstrated that TP53 activity suppresses the mevalonate pathway [10], which may explain the increased mevalonate pathway. In both FulvR and TamR cells, we identified a significant decrease in TP53 binding protein 1 (TP53BP1) after DMF treatment (**SFigure 10**). When bound to TP53, TP53BP1 stabilizes and enhances the activity of TP53 [11, 12]. Our observed downregulation of TP53BP1 suggests decreased TP53 activity, which would relieve TP53’s inhibitory effect on the mevalonate pathway, thereby promoting the increased mevalonate pathway enzyme expression seen in FulvR and TamR cells. Together, our data suggest that DMF reprograms lipid metabolism in Fulv and Tam resistance, potentially by decreasing TP53 activity and enhancing steroid hormone production through upregulated cholesterol synthesis. Next, we focused on how Tam and Fulv drug resistance and the subsequent reversal of resistance by DMF impacted estrogen receptor DNA binding activity.

### DMF treatment restores ER binding and transcriptional activity in TamR cells

To investigate whether DMF reversed endocrine resistance in TamR cells by modifying ER DNA binding activity, we performed CUT&RUN analysis. We compared ER binding profiles across TamR and parental cells under different treatment conditions. We combined the sequenced datasets from MCF7 and UCD12 cells to consider biological variability. In the vehicle-treated (0.1% ethanol) parental cells, we detected 4,926 total peaks, compared to 1,871 detected peaks in vehicle-treated TamR cells. A total of 713 peaks (11.7%) were common to parental and TamR vehicle-treated cells (**Figure 6A**). TamR cells treated with DMF had 2,933 identified peaks, 1,106 (16.4%) of which were shared with vehicle-treated ER peaks, suggesting partial restoration of ER binding activity. These results indicated that DMF treatment in TamR cells enhances ER’s ability to bind DNA similarly to parental cells.

**Figure 6.**
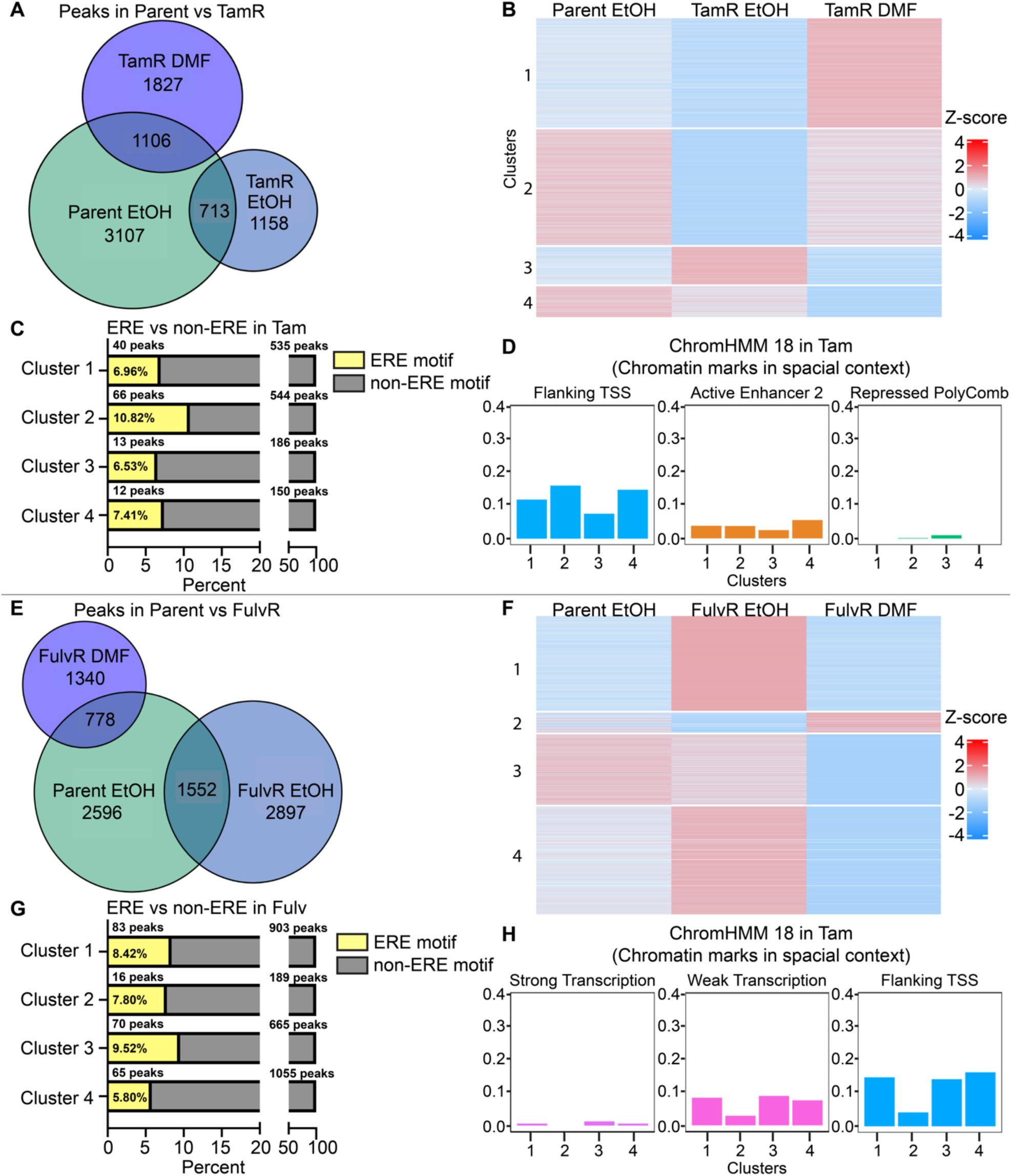
Estrogen receptor DNA binding in DMF reversal of Tam and Fulv resistance are differentially regulated. Tam sensitive and TamR cells were treated with vehicle (EtOH) or 50μM DMF. **(A)** Venn diagram of total number of identified peaks after CUT&RUN for ERalpha. **(B)** Z-score k-means peak clustering of Cut&Run scores. **(C)** Percentage of peaks with ERE motifs vs non-ERE motifs. **(D)** Chromatin marking by clusters. **(E-H)** Same as A-D but for Fulv sensitive and FulvR cells treated with vehicle (EtOH) or 50μM DMF.

We used HOMER motif analysis to identify known ER binding motifs and assess whether DMF treatment contributed to reversing Tam resistance by altering ER’s DNA-binding behavior. Specifically, we compared ER binding motifs in DMF-treated TamR cells and vehicle-treated TamR cells to those in vehicle-treated parental (Tam-sensitive) cells. Once again, our goal was to determine if DMF treatment caused ER binding patterns in resistant TamR cells to revert to a state more similar to the parental cells responsive to Tam. HOMER motif analysis identified 239 common ER binding motifs between DMF-treated TamR cells and parental cells, compared to 185 motifs shared between vehicle-treated TamR cells and parental cells. This supports the conclusion that DMF treatment in TamR cells partially restores a parental ER binding pattern (**SFigure 11A**).

To visualize the binding of ER to DNA under different treatment conditions, we used k-means clustering of the spiked-in normalized CUT&RUN scores and generated a heatmap (**Figure 6B**). Vehicle-treated parental cells had strong binding activity in clusters 2 and 4, while TamR cells treated with vehicle had strong binding activity in cluster 3, decreased activity in cluster 4, and an absence of activity in cluster 2. This binding pattern showed that Tam resistance resulted in the loss of ER binding activity associated with cluster 2. However, after DMF treatment, TamR cells gained a strong ER binding activity in cluster 1, and regained ER binding in cluster 2. Therefore, cluster 2 best represents the portion of ER binding that is reverted to parental patterns in DMF-treated TamR cells. Additionally, we found that cluster 2 had the largest percentage of peaks binding to EREs (approximately 11%, **Figure 6C**).

Finally, we analyzed the chromatin states associated with ER binding clusters using ChromHMM annotations [13]. Cluster 2, the cluster presenting strong binding in vehicle-treated parental cells and was associated with DMF-treated TamR cells, showed increased marks of active transcription (TssFlnk, TxWk) and enhancers (EnhA2, EnhWk), indicating a restoration of transcriptional activity near ER binding sites (**Figure 6D and SFigure 11B**). Recalling that DMF alone decreased TamR cell growth, but a combination treatment of Tam plus DMF further reduced TamR growth, we conclude that the peaks represented in Cluster 2 are the peaks that revert to parental patterns and are susceptible to Tam treatment. This contrasted with the repressive states observed in clusters defining vehicle-treated TamR cells, especially the increased repressor polycomb mark (ReprPC) in cluster 3 (**Figure 6D and SFigure 11B**). Together, these data reinforced the role of DMF in reactivating ER-driven gene expression.

Overall, our data suggest that ER DNA binding activity in TamR cells differs from that of parental cells in terms of the number of common peaks, binding of EREs, and location on the chromatin. However, DMF treatment reverts ER binding activity to a similar pattern as the parental Tam-sensitive cells.

### DMF treatment disrupts ER binding and alters regulatory networks in FulvR cells

Unlike TamR cells, FulvR cells showed a different pattern of ER DNA binding after DMF treatment. Vehicle-treated FulvR cells had 4,449 ER binding peaks, with 1,552 peaks (31.5%) shared with parental cells. Following DMF treatment, the total number of ER binding peaks in FulvR cells was nearly halved to 2,118, with only 778 peaks (15.7%) overlapping with parental cells (**Figure 6E**). This indicated that DMF treatment significantly altered the ER binding landscape differently in FulvR cells compared to TamR cells. HOMER motif analysis further revealed that ER binding in DMF-treated FulvR cells diverged from both vehicle-treated FulvR and parental cells, with only 180 shared motifs compared to 288 between vehicle-treated groups (**SFigure 12A**). This suggests that the ER binding activity in DMF-treated FulvR cells is distinct from that of vehicle-treated parent (Fulv sensitive) cells.

Clustering analysis highlighted large differences in ER binding patterns between DMF-treated FulvR cells and all other groups (**Figure 6F**). Vehicle-treated parent cells (Fulv sensitive) were strongly defined by cluster 3 and weakly by cluster 4. The percentage of peaks binding ERE motifs was highest in cluster 3 (**Figure 6G**). Vehicle-treated FulvR cells (Fulv resistant) were strongly defined by clusters 1 and 4 and weakly by cluster 3. These groups were in stark contrast to DMF-treated FulvR cells, which regained sensitivity to Fulv but were mainly defined by cluster 2. Intersecting peaks from cluster 2 with ChromHMM-18 annotations demonstrated that this cluster lacked transcriptionally active states. Specifically, strong transcription (Tx), weak transcription (TxWk), and flanking transcriptional start sites (TssFlnk) were absent in cluster 2 (**Figure 6H** and **SFigure 12B**). This suggests that DMF significantly alters the regulatory interactions of ER in FulvR cells, possibly diminishing ER’s capacity to maintain transcriptional programs associated with proliferation.

To identify if there were TF motifs that might cooperate with ER and help explain the transcription chromosome states, we input the TF motifs from HOMER into a FIMO analysis, which allowed us to identify what fraction of the motifs were represented in our samples. We identified TF motifs in vehicle-treated FulvR cells with a fraction cutoff of 0.10 or higher. Next, we determined the fold change between the TF motif fraction of vehicle-treated FulvR cells and DMF-treated FulvR cells. We identified 17 TF motifs in FulvR vehicle-treated cells with greater than 1.6 fold-change fraction compared to DMF-treated FulvR cells, 7 of which have been identified as having a cooperative interaction with ER or having the ability to affect endocrine resistance (**Table 1**). DMF treatment decreased the fractions of TF representation compared to vehicle-treated, suggesting that FulvR cells are not entirely independent of cooperative interactions between ER and that other ER-modifying TFs could affect FulvR cell proliferation.

**Table 1.**
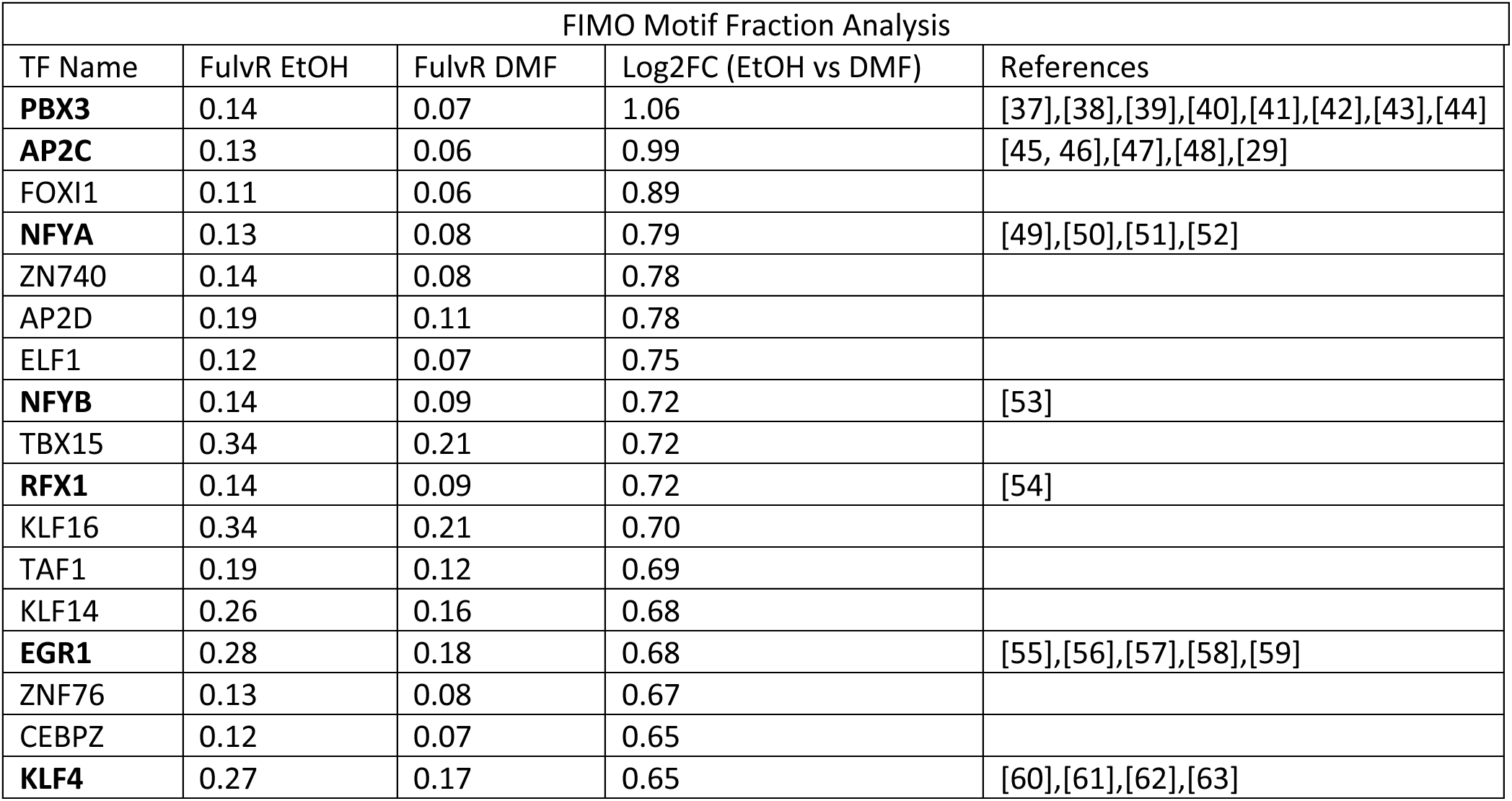
Top-ranked transcription factors from FIMO analysis that interact with ER. The transcription factors in bold type are known to interact with ER or influence endocrine therapy response.

## DISCUSSION

The development of endocrine resistance in ER+ breast cancer remains a significant hurdle, often leading to disease progression and limited treatment options [14, 15]. Endocrine-resistant breast cancer cells are known to undergo substantial metabolic reprogramming [16, 17]. In this study, we examined commonalities and divergence in the development of Tam or Fulv resistance by generating multiple endocrine-resistant ER+ breast cancer cell lines. Our results were consistent with the emerging landscape of acquired endocrine resistance and showed a substantial role for metabolic reprogramming through alterations in nucleotide metabolism, amino acid utilization, and redox balance. Many of these disruptions are known to provide tumor cells with growth and survival advantages [18, 19], which is one reason targeting cancer metabolism has emerged as a promising approach to combat drug resistance [15].

Our data also pointed to the TCA cycle as a potential therapeutic target, so we investigated the use of DMF to overcome this metabolic blockade observed in both Tam and FulvR cells. DMF is a cell-permeable ester that is metabolized to fumarate, replenishing TCA cycle intermediates [4]. Clinically approved for treating multiple sclerosis and psoriasis due to its immunomodulatory and cytoprotective effects [20, 21], DMF has also demonstrated inhibitory effects on ER+ breast cancer cells by targeting nuclear factor kappa-light-chain-enhancer of activated B cells (**NFkB**) on the growth of MCF7 ER+ breast cancer cells [22], and via inhibition of zinc finger protein 217 (**ZNF217**) [23]. Notably, overexpression of ZNF217 in MCF-7 cells has been associated with Tam resistance [24]. Our data demonstrated that DMF treatment effectively restored the TCA cycle blockade in FulvR cells and reversed both Fulv and Tam endocrine resistance, although through different mechanisms.

Tam is a known partial agonist of the estrogen receptor and can weakly activate its ability to stimulate cell proliferation pathways as Tam resistance develops [25]. We show that DMF treatment reduced cell proliferation in TamR cells and restored ER DNA binding activity to a pattern resembling that of parental, endocrine-sensitive cells. Notably, a similar ER binding pattern was associated with the ChromHMM-18 markers, indicative of active transcription. This finding suggests a direct link between metabolic state and ER function, indicating that metabolic correction can reinstate normal ER signaling pathways disrupted during the development of Tam resistance.

Reversal of Fulv and Tam resistance was associated with decreased activation of the mevalonate pathway. The mevalonate pathway is crucial for cholesterol synthesis and the prenylation of proteins involved in cell proliferation and survival [26]. Activation of the mevalonate pathway in DMF-treated FulvR and TamR cells may have been due to decreased TP53 activity associated with reduced expression of its co-regulator TP53BP1 [10]. Interestingly, vehicle-treated TamR cells had significantly decreased citrate levels, and vehicle-treated FulvR cells had significantly decreased 2-oxoglutarate levels, both of which were restored with DMF treatment and both of which can be used to fuel acetyl-CoA production to sustain elevated cholesterol synthesis [27]. These findings suggest that the mevalonate pathway is an important mediator of DMF-mediated reversal of Tam and Fulv resistance.

Metabolic reprogramming contributed to ER DNA binding activity differently in Tam and Fulv resistance. Unlike TamR cells, DMF-treated FulvR cells did not regain the parental ER DNA binding signature, which is consistent with fulvestrant’s mechanism of action – inducing degradation of ER [28]. However, our FIMO discovery analysis suggested that DMF treatment in FulvR cells led to particular ER/TF cooperative interactions to reduce FulvR proliferation (**STable 1** and [29, 30]).

Currently, the most successful compounds targeting metabolism in breast cancer are inhibitors of the PI3K/Akt/mTOR pathway. Drugs like everolimus, alpelisib, and capivasertib are indicated for use in combination with fulvestrant in patients who have a targeted mutation and have progressed on an endocrine-based regimen [31–33]. Everolimus is used in the setting of recurrent or metastatic ER+ breast cancer [34]. Alpelisib is prescribed for patients with a

PIK3CA mutation [35], and capivasertib is used in patients with a mutation in PIK3CA or AKT1 or who have a PTEN loss [36]. These drugs are particularly effective when combined with endocrine therapies such as aromatase inhibitors or fulvestrant, highlighting the interconnected nature of ER and the PI3K/Akt/mTOR metabolic pathway. However, questions remain about using these or new compounds in tumors that do not express these mutations. Therefore, understanding the reprogrammed metabolic networks in cancer cells and identifying novel therapeutic targets remain critical areas of ongoing research. Our results highlight the importance of metabolic adaptations on ER DNA binding in breast cancer and demonstrate that targeting adaptations can restore ER activity and therapeutic sensitivity. Our study strengthens the current rationale for integrating metabolic modulators like DMF into combination therapy regimens to improve outcomes in patients who develop endocrine-resistant disease.

## Conflict of interest

The authors have declared that no conflict of interest exists.

## Funding

This work was supported by the National Institutes of Health R01CA205044 (P. Kabos), NIH grant P20CA046934 supporting the University of Colorado Cancer Center Genomics Shared Resource, the University of Colorado Mass Spectrometry Metabolomics Shared Resource Facility, and the Mass Spectrometry Proteomics Shared Resource.

## Notes

### Competing Interest Statement

The authors have declared no competing interest.

## REFERENCES

1. Skene, P.J., J.G. Henikoff, and S. Henikoff, Targeted in situ genome-wide profiling with high efficiency for low cell numbers. Nature Protocols, 2018. 13(5): p. 1006–1019.

2. Han, A.L., et al., Estradiol (E2) concentration shapes the chromatin binding landscape of the estrogen receptor. 2022, Cold Spring Harbor Laboratory.

3. Lloyd, S., Least squares quantization in PCM. IEEE Transactions on Information Theory, 1982. 28(2): p. 129–137.

4. Kourakis, S., et al., Dimethyl Fumarate and Its Esters: A Drug with Broad Clinical Utility? Pharmaceuticals (Basel), 2020. 13(10).

5. Bhargava, P., et al., Dimethyl fumarate treatment induces lipid metabolism alterations that are linked to immunological changes. Ann Clin Transl Neurol, 2019. 6(1): p. 33–45.

6. Blumenfeld Kan, S., et al., HDL-cholesterol elevation associated with fingolimod and dimethyl fumarate therapies in multiple sclerosis. Mult Scler J Exp Transl Clin, 2019. 5(4): p. 2055217319882720.

7. Borgquist, S., et al., Prognostic impact of tumour-specific HMG-CoA reductase expression in primary breast cancer. Breast Cancer Res, 2008. 10(5): p. R79.

8. Scott, O.W., et al., Post-diagnostic statin use and breast cancer-specific mortality: a population-based cohort study. Breast Cancer Res Treat, 2023. 199(1): p. 195–206.

9. Smith, A., et al., Pre-diagnostic statin use, lymph node status and mortality in women with stages I-III breast cancer. Br J Cancer, 2017. 117(4): p. 588–596.

10. Moon, S.H., et al., p53 Represses the Mevalonate Pathway to Mediate Tumor Suppression. Cell, 2019. 176(3): p. 564–580 e19.

11. Cuella-Martin, R., et al., 53BP1 Integrates DNA Repair and p53-Dependent Cell Fate Decisions via Distinct Mechanisms. Mol Cell, 2016. 64(1): p. 51–64.

12. Belal, H., E.F. Ying Ng, and F. Meitinger, 53BP1-mediated activation of the tumor suppressor p53. Curr Opin Cell Biol, 2024. 91: p. 102424.

13. Ernst, J. and M. Kellis, ChromHMM: automating chromatin-state discovery and characterization. Nat Methods, 2012. 9(3): p. 215–6.

14. Musgrove, E.A. and R.L. Sutherland, Biological determinants of endocrine resistance in breast cancer. Nat Rev Cancer, 2009. 9(9): p. 631–43.

15. Osborne, C.K. and R. Schiff, Mechanisms of endocrine resistance in breast cancer. Annu Rev Med, 2011. 62: p. 233–47.

16. Martinez-Outschoorn, U.E., et al., Cancer metabolism: a therapeutic perspective. Nat Rev Clin Oncol, 2017. 14(1): p. 11–31.

17. Pavlova, N.N., J. Zhu, and C.B. Thompson, The hallmarks of cancer metabolism: Still emerging. Cell Metab, 2022. 34(3): p. 355–377.

18. DeBerardinis, R.J., et al., The biology of cancer: metabolic reprogramming fuels cell growth and proliferation. Cell Metab, 2008. 7(1): p. 11–20.

19. Pavlova, N.N. and C.B. Thompson, The Emerging Hallmarks of Cancer Metabolism. Cell Metab, 2016. 23(1): p. 27–47.

20. Gold, R., et al., Placebo-controlled phase 3 study of oral BG-12 for relapsing multiple sclerosis. N Engl J Med, 2012. 367(12): p. 1098–107.

21. Longbrake, E.E., et al., Dimethyl fumarate treatment shifts the immune environment toward an anti-inflammatory cell profile while maintaining protective humoral immunity. Mult Scler, 2021. 27(6): p. 883–894.

22. Kastrati, I., et al., Dimethyl Fumarate Inhibits the Nuclear Factor κB Pathway in Breast Cancer Cells by Covalent Modification of p65 Protein. Journal of Biological Chemistry, 2016. 291(7): p. 3639–3647.

23. Sharma, T., et al., Dimethyl fumarate inhibits ZNF217 and can be beneficial in a subset of estrogen receptor positive breast cancers. Breast Cancer Res Treat, 2023. 201(3): p. 561–570.

24. Nguyen, N.T., et al., A functional interplay between ZNF217 and estrogen receptor alpha exists in luminal breast cancers. Mol Oncol, 2014. 8(8): p. 1441–57.

25. MacNab, M.W., R.J. Tallarida, and R. Joseph, An evaluation of tamoxifen as a partial agonist by classical receptor theory--an explanation of the dual action of tamoxifen. Eur J Pharmacol, 1984. 103(3-4): p. 321–6.

26. Mullen, P.J., et al., The interplay between cell signalling and the mevalonate pathway in cancer. Nat Rev Cancer, 2016. 16(11): p. 718–731.

27. Mullen, A.R., et al., Reductive carboxylation supports growth in tumour cells with defective mitochondria. Nature, 2011. 481(7381): p. 385–8.

28. Robertson, J.F., Fulvestrant (Faslodex) -- how to make a good drug better. Oncologist, 2007. 12(7): p. 774–84.

29. Woodfield, G.W., et al., TFAP2C controls hormone response in breast cancer cells through multiple pathways of estrogen signaling. Cancer Res, 2007. 67(18): p. 8439–43.

30. Tan, S.K., et al., AP-2gamma regulates oestrogen receptor-mediated long-range chromatin interaction and gene transcription. EMBO J, 2011. 30(13): p. 2569–81.

31. Cerma, K., et al., Targeting PI3K/AKT/mTOR Pathway in Breast Cancer: From Biology to Clinical Challenges. Biomedicines, 2023. 11(1).

32. Jones, R.H., et al., Fulvestrant plus capivasertib versus placebo after relapse or progression on an aromatase inhibitor in metastatic, oestrogen receptor-positive breast cancer (FAKTION): a multicentre, randomised, controlled, phase 2 trial. Lancet Oncol, 2020. 21(3): p. 345–357.

33. Owonikoko, T.K. and F.R. Khuri, Targeting the PI3K/AKT/mTOR pathway: biomarkers of success and tribulation. Am Soc Clin Oncol Educ Book, 2013.

34. Royce, M.E. and D. Osman, Everolimus in the Treatment of Metastatic Breast Cancer. Breast Cancer (Auckl), 2015. 9: p. 73–9.

35. Andre, F., et al., Alpelisib for PIK3CA-Mutated, Hormone Receptor-Positive Advanced Breast Cancer. N Engl J Med, 2019. 380(20): p. 1929–1940.

36. Nierengarten, M.B., FDA approves capivasertib with fulvestrant for breast cancer. Cancer, 2024. 130(6): p. 835–836.

37. Kao, T.W., et al., PBX1 as a novel master regulator in cancer: Its regulation, molecular biology, and therapeutic applications. Biochim Biophys Acta Rev Cancer, 2024. 1879(2): p. 189085.

38. Pavithran, H. and R. Kumavath, Emerging role of pioneer transcription factors in targeted ERα positive breast cancer. Explor Target Antitumor Ther, 2021. 2(1): p. 26–35.

39. Pang, Z.-y., et al., Leptin-elicited PBX3 confers letrozole resistance in breast cancer. Endocrine-Related Cancer, 2021. 28(3): p. 173–189.

40. Liu, Y., et al., The regulation of PBXs and their emerging role in cancer. J Cell Mol Med, 2022. 26(5): p. 1363–1379.

41. Ao, X., et al., PBX1 is a valuable prognostic biomarker for patients with breast cancer. Exp Ther Med, 2020. 20(1): p. 385–394.

42. Magnani, L., et al., PBX1 genomic pioneer function drives ERalpha signaling underlying progression in breast cancer. PLoS Genet, 2011. 7(11): p. e1002368.

43. Yang, H., et al., A network-based approach reveals the dysregulated transcriptional regulation in non-alcoholic fatty liver disease. iScience, 2021. 24(11): p. 103222.

44. Eeckhoute, J., et al., Positive cross-regulatory loop ties GATA-3 to estrogen receptor alpha expression in breast cancer. Cancer Res, 2007. 67(13): p. 6477–83.

45. Franke, C.M., et al., TFAP2C regulates carbonic anhydrase XII in human breast cancer. Oncogene, 2020. 39(6): p. 1290–1301.

46. Yuan, L., et al., TFAP2C Activates CST1 Transcription to Facilitate Breast Cancer Progression and Suppress Ferroptosis. Biochem Genet, 2024. 62(5): p. 3858–3875.

47. Woodfield, G.W., et al., Interaction of TFAP2C with the estrogen receptor-alpha promoter is controlled by chromatin structure. Clin Cancer Res, 2009. 15(11): p. 3672–9.

48. Cyr, A.R., et al., TFAP2C governs the luminal epithelial phenotype in mammary development and carcinogenesis. Oncogene, 2015. 34(4): p. 436–44.

49. De Amicis, F., et al., Resveratrol, through NF-Y/p53/Sin3/HDAC1 complex phosphorylation, inhibits estrogen receptor alpha gene expression via p38MAPK/CK2 signaling in human breast cancer cells. FASEB J, 2011. 25(10): p. 3695–707.

50. Ying, S., et al., Estrogen receptor alpha and nuclear factor Y coordinately regulate the transcription of the SUMO-conjugating UBC9 gene in MCF-7 breast cancer cells. PLoS One, 2013. 8(9): p. e75695.

51. Farsetti, A., et al., Inhibition of ERα-Mediated Trans-Activation of Human Coagulation Factor XII Gene by Heteromeric Transcription Factor NF-Y. Endocrinology, 2001. 142(8): p. 3380–3388.

52. Kang, K., et al., Predicting FOXM1-Mediated Gene Regulation through the Analysis of Genome-Wide FOXM1 Binding Sites in MCF-7, K562, SK-N-SH, GM12878 and ECC-1 Cell Lines. Int J Mol Sci, 2020. 21(17).

53. Notas, G., et al., Whole transcriptome analysis of the ERalpha synthetic fragment P295-T311 (ERalpha17p) identifies specific ERalpha-isoform (ERalpha, ERalpha36)-dependent and - independent actions in breast cancer cells. Mol Oncol, 2013. 7(3): p. 595–610.

54. Shibata, M., et al., Expression of regulatory factor X1 can predict the prognosis of breast cancer. Oncol Lett, 2017. 13(6): p. 4334–4340.

55. Shajahan-Haq, A.N., et al., EGR1 regulates cellular metabolism and survival in endocrine resistant breast cancer. Oncotarget, 2017. 8(57): p. 96865–96884.

56. Horibata, S., et al., A bi-stable feedback loop between GDNF, EGR1, and ERalpha contribute to endocrine resistant breast cancer. PLoS One, 2018. 13(4): p. e0194522.

57. Yan, S., et al., Targeting the crosstalk between estrogen receptors and membrane growth factor receptors in breast cancer treatment: Advances and opportunities. Biomedicine & Pharmacotherapy, 2024. 175: p. 116615.

58. Kim, H.R., et al., Egr1 is rapidly and transiently induced by estrogen and bisphenol A via activation of nuclear estrogen receptor-dependent ERK1/2 pathway in the uterus. Reprod Toxicol, 2014. 50: p. 60–7.

59. Marks, B.A., et al., GDNF-RET signaling and EGR1 form a positive feedback loop that promotes tamoxifen resistance via cyclin D1. BMC Cancer, 2023. 23(1): p. 138.

60. Hu, D., et al., Novel insight into KLF4 proteolytic regulation in estrogen receptor signaling and breast carcinogenesis. J Biol Chem, 2012. 287(17): p. 13584–97.

61. Jia, Y., et al., KLF4 overcomes tamoxifen resistance by suppressing MAPK signaling pathway and predicts good prognosis in breast cancer. Cell Signal, 2018. 42: p. 165–175.

62. Simmen, R.C., et al., The emerging role of Krüppel-like factors in endocrine-responsive cancers of female reproductive tissues. J Endocrinol, 2010. 204(3): p. 223–31.

63. Zhou, Z., et al., Regulation of KLF4 by posttranslational modification circuitry in endocrine resistance. Cell Signal, 2020. 70: p. 109574.

