## Supplemental Material for "Metabolic Switch in Endocrine Resistant Estrogen Receptor Positive Breast Cancer"

Conflict of interest: The authors have declared that no conflict of interest exists.

Corresponding Author Information

Peter Kabos
University of Colorado Denver AMC
12801 E 17th Ave
RC1 South, M.S. 8117
Aurora CO 80045
303-724-4690
ORCID: 0000-0003-1500-471X

Heather Brechbuhl
University of Colorado Denver AMC
12801 E 17th Ave
RC1 South, M.S. 8117
Aurora CO 80045
303-724-5823
ORCID: 0000-0003-3132-4465

Figures: 6 main; 12 supplemental

Tables: 1 main

**KEYWORDS**

Breast Cancer, Estrogen Receptor, Endocrine Resistance, Metabolism.

Funding: This work was supported by the National Institutes of Health R01CA205044 (P. Kabos), NIH grant P20CA046934 supporting the University of Colorado Cancer Center Genomics Shared Resource, the University of Colorado Mass Spectrometry Metabolomics Shared Resource Facility, and the Mass Spectrometry Proteomics Shared Resource.

Star Methods

| REAGENT or RESOURCE | SOURCE | IDENTIFIER |
|---|---|---|
| Antibodies | | |
| Estrogen Receptor alpha polyclonal | Abcam | Cat# Ab3575<br>RRID:AB_303921 |
| Bacterial and virus strains | | |
| N/A | | |
| Biological samples | | |
| N/A | | |
| Chemicals, peptides, and recombinant proteins | | |
| Fulvestrant (ICI-182780) | Selleckchem.com | Cat# S1191 |
| Tamoxifen | Cayman Chemical | Cat# 13258 |
| DMEM | Corning | Cat# 10-013-CV |
| DMEM/F-12 | Corning | Cat# 10-092-CV |
| RPMI 1640 | Corning | Cat# 10-040-CV |
| Insulin | Sigma Chemicals | Cat# I-6634 |
| Equalfetal Bovine Serum, Alternative to FBS, US Origin | Atlas Biologicals | Cat# EF-0500-A |
| Critical commercial assays | | |
| Universal Mycoplasma Detection Kit | ATCC | Cat# 30-1012K |
| Pierce™ Detergent Compatible Bradford assay | Thermo Fisher Scientific | Cat# 1863028 |
| TruSeq® Stranded mRNA Library Prep | Illumina | Cat# 20020595 |

| | | |
|---|---|---|
| Deposited data | | |
| Experimental models: Cell lines | | |
| BT474 parent is from ATCC but passage matched to the TamR and FulvR cell lines | This paper | ATCC HTB-20 |
| MCF7 parent is from ATCC but passage matched to the TamR and FulvR cell lines | This paper | ATCC HTB-22 |
| MDA-MB-134-VI parent is from ATCC but passage matched to the TamR and FulvR cell lines | This paper | ATCC HTB-23 |
| T47D parent is from ATCC but passage matched to the TamR and FulvR cell lines | This paper | ATCC HTB-133 |
| UCD12 parent is a PDX generated at the University of Colorado and is passage matched to the TamR and FulvR cell lines | University of Colorado ER+ Breast Tissue Bank | Finlay-Schultz, J. et al., 2020 [1] |
| ZR75-1 parent is from ATCC but passage matched to the TamR and FulvR cell lines | This paper | ATCC CRL-1500 |
| BT474 TamR | This paper | |
| BT474 FulvR | This paper | |
| MCF7 TamR | This paper | |
| MCF7 FulvR | This paper | |
| MDA-MB-134-VI TamR | This paper | |
| MDA-MB-134-VI FulvR | This paper | |
| T47D TamR | This paper | |
| T47D FulvR | This paper | |
| UCD12 TamR | This paper | |
| UCD12 FulvR | This paper | |
| ZR75-1 TamR | This paper | |
| ZR75-1 FulvR | This paper | |

| | | |
|---|---|---|
| Experimental models: Organisms/strains | | |
| N/A | | |
| Oligonucleotides | | |
| N/A | | |
| Recombinant DNA | | |
| N/A | | |
| Software and algorithms | | |
| Cutadapt 4.1 | https://cutadapt.readthedocs.io/en/stable | Martin M., 2011 [2] |
| STAR | https://github.com/alexdobin/STAR | Dobin, A. et al., 2013 [3] |
| DESEQ2 | https://github.com/mikelove/DESeq2 | Love, M. et al., 2014 [4] |
| Bowtie2 2.3.4.3 | http://bowtie-bio.sourceforge.net/bowtie2/index.shtml | Langmed, B. et al., 2012 [5] |
| Samtools | http://samtools.sourceforge.net/ | Li, H. et al., 2009 [6] and<br>Li, H., 2011 [7] |
| Python 3.6.5 | https://www.python.org/ | Python Software Foundation |
| R 4.0.3 | https://www.r-project.org | The R Foundation |
| Perl 5.26 | https://www.perl.org | The Perl Foundation |
| K means clustering | | Lloyd, S. 1982 [8] |
| FIMO | https://meme-suite.org/meme/doc/fimo.html | Bailey TL., et al. 2015 [9]<br>and<br>Grant CE., et al. 2011 [10] |

| | | |
|---|---|---|
| MetaboAnalyst | https://www.metaboanalyst.ca/MetaboAnalyst/home.xhtml | Xia, J. et al., 2009 [11] |
| GraphPad Prism Version 9.5.1 | www.graphpad.com | GraphPad by Dotmatics |
| Other | | |

**Supplemental Figure 1. <u>Joint Pathway Analysis in MCF7 Parent vs FulvR or Parent vs TamR</u>**. All curated pathways identified by JPA with have a p <0.019, false discovery rate (FDR) < 0.055, and a composite score > 1.0.

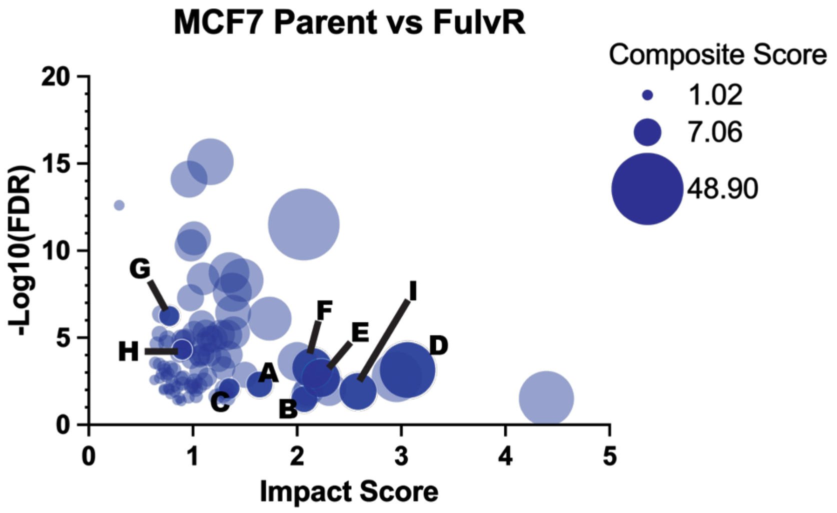

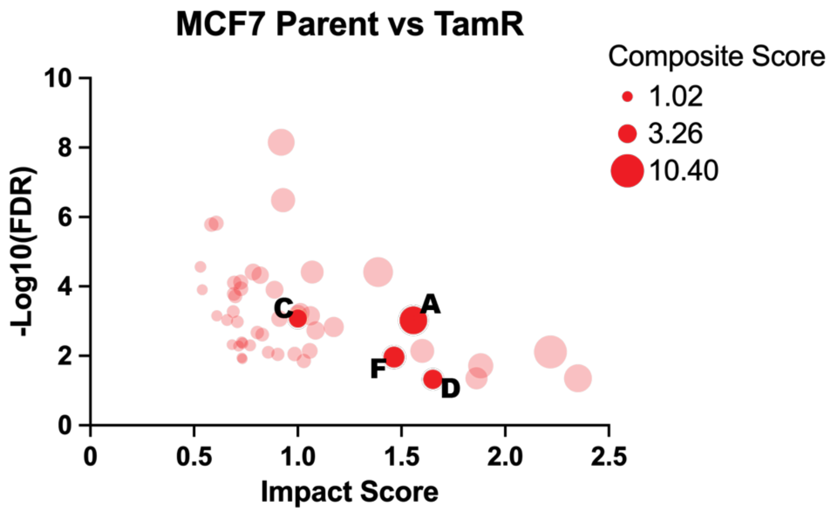

**Key**

**A. Alanine, aspartate and glutamate metabolism**
**B. Citrate cycle (TCA cycle)**
**C. Cysteine and methionine metabolism**
**D. EGFR tyrosine kinase inhibitor resistance**
**E. ErbB signaling pathway**
**F. Glutathione metabolism**
**G. mTOR signaling pathway**
**H. PI3K-Akt signaling pathway**
**I. Pyruvate metabolism**

**Supplemental Figure 2.** **Joint Pathway Analysis in T47D Parent vs FulvR or Parent vs TamR**. All curated pathways identified by JPA with have a p <0.019, false discovery rate (FDR) < 0.055, and a composite score > 1.0.

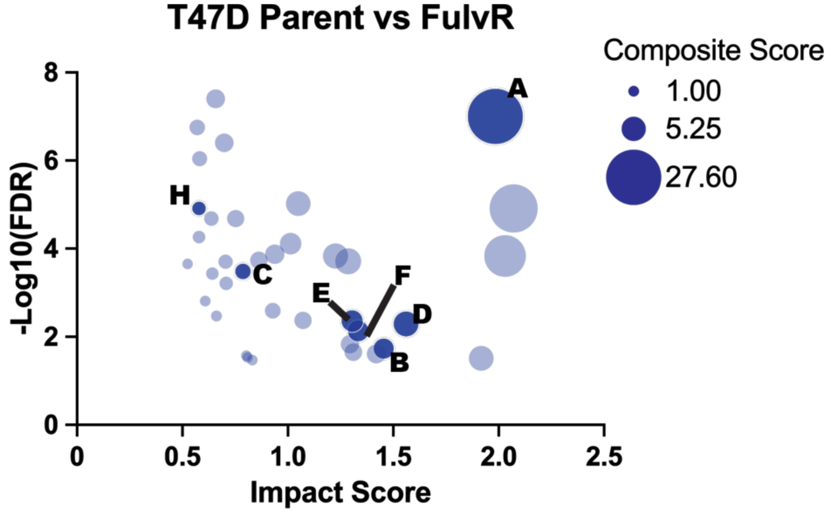

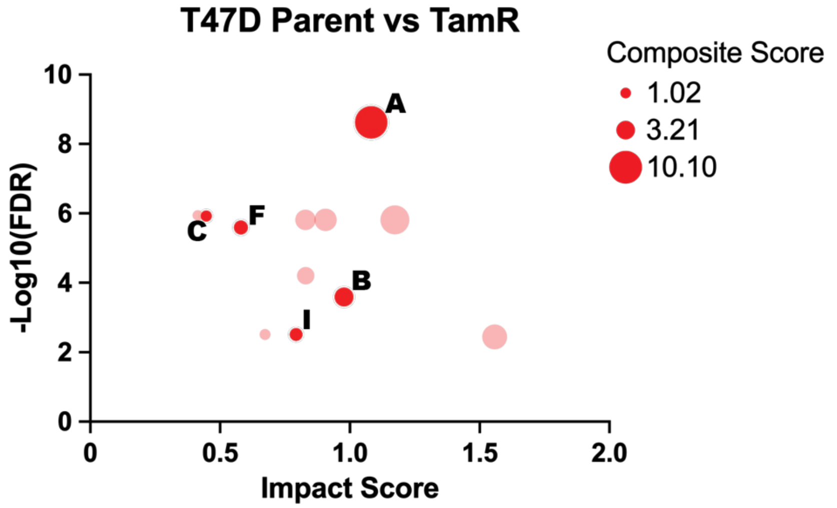

**Key**

**A. Alanine, aspartate and glutamate metabolism**
**B. Citrate cycle (TCA cycle)**
**C. Cysteine and methionine metabolism**
**D. EGFR tyrosine kinase inhibitor resistance**
**E. ErbB signaling pathway**
**F. Glutathione metabolism**
**G. mTOR signaling pathway**
**H. PI3K-Akt signaling pathway**
**I. Pyruvate metabolism**

**Supplemental Figure 3.** **Joint Pathway Analysis in ZR75 Parent vs FulvR or Parent vs TamR**. All curated pathways identified by JPA with have a p <0.019, false discovery rate (FDR) < 0.055, and a composite score > 1.0.

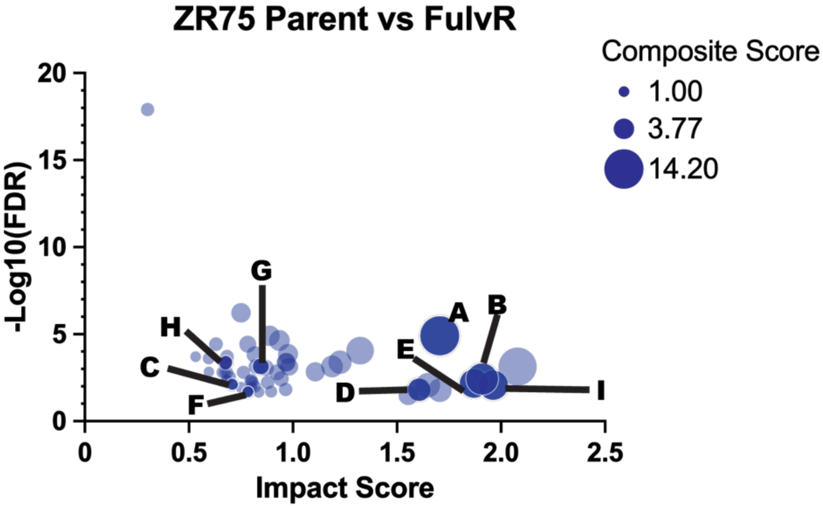

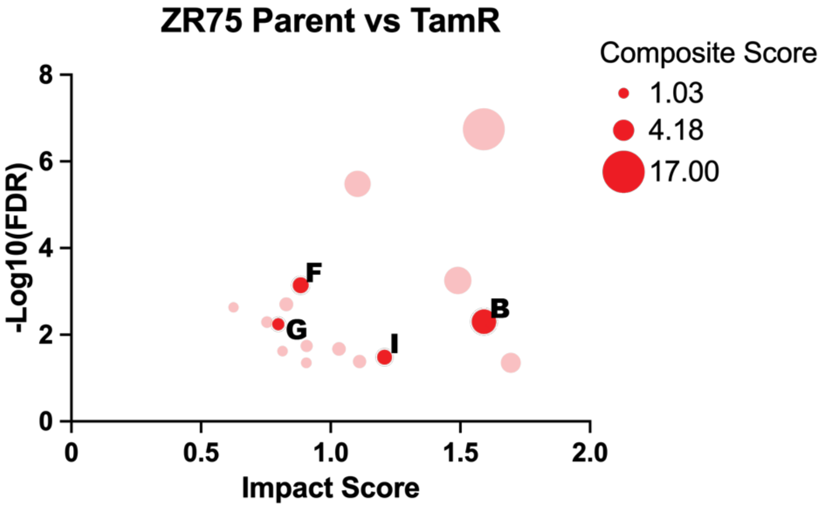

**Key**

**A. Alanine, aspartate and glutamate metabolism**
**B. Citrate cycle (TCA cycle)**
**C. Cysteine and methionine metabolism**
**D. EGFR tyrosine kinase inhibitor resistance**
**E. ErbB signaling pathway**
**F. Glutathione metabolism**
**G. mTOR signaling pathway**
**H. PI3K-Akt signaling pathway**
**I. Pyruvate metabolism**

**Supplemental Figure 4. Joint Pathway Analysis in UCD12 Parent vs FulvR or Parent vs TamR**. All curated pathways identified by JPA with have a p <0.019, false discovery rate (FDR) < 0.055, and a composite score > 1.0.

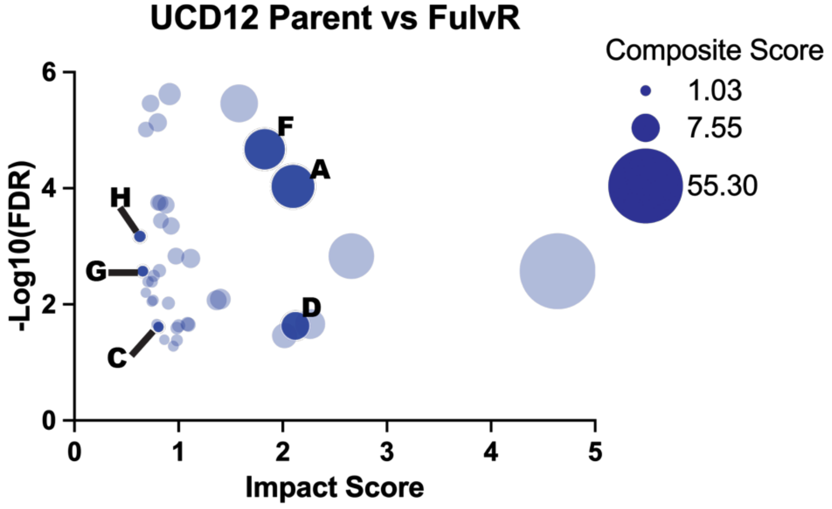

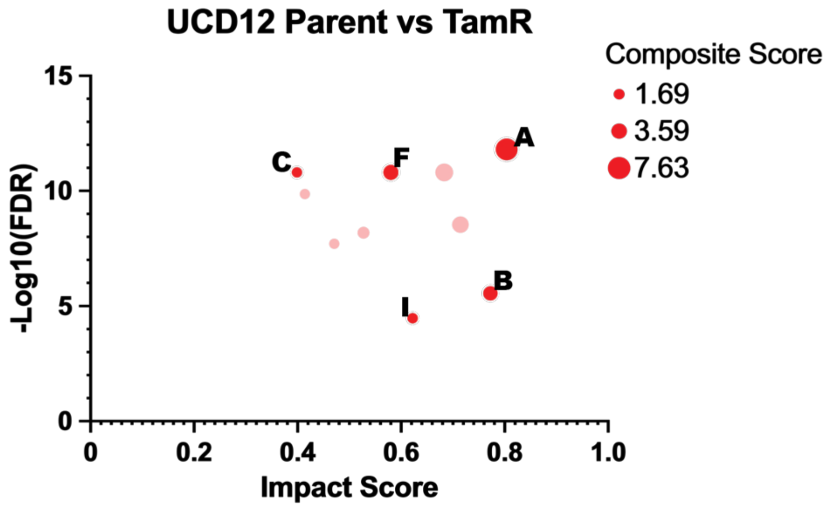

**Key**

**A. Alanine, aspartate and glutamate metabolism**
**B. Citrate cycle (TCA cycle)**
**C. Cysteine and methionine metabolism**
**D. EGFR tyrosine kinase inhibitor resistance**
**E. ErbB signaling pathway**
**F. Glutathione metabolism**
**G. mTOR signaling pathway**
**H. PI3K-Akt signaling pathway**
**I. Pyruvate metabolism**

**Supplemental Figure 5.** **<u>Joint Pathway Analysis in BT474 Parent vs FulvR or Parent vs TamR</u>**. All curated pathways identified by JPA with have a p <0.019, false discovery rate (FDR) < 0.055, and a composite score > 1.0.

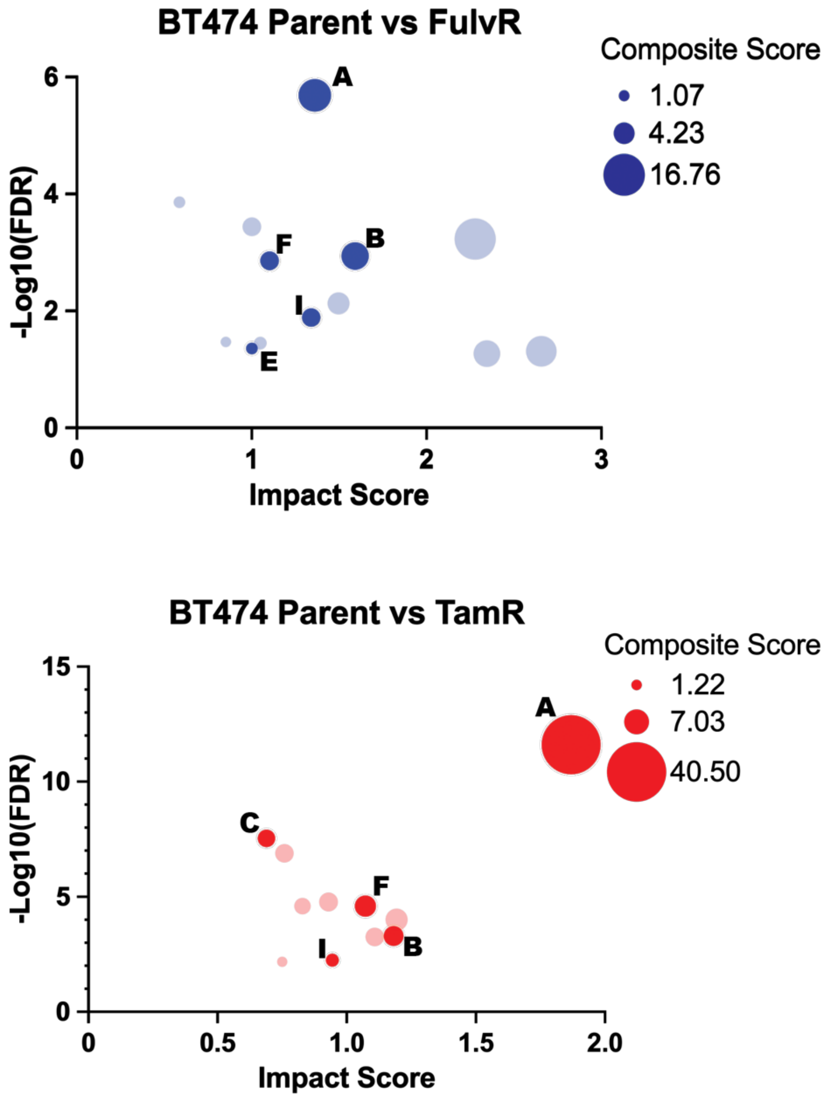

**Key**

**A. Alanine, aspartate and glutamate metabolism**
**B. Citrate cycle (TCA cycle)**
**C. Cysteine and methionine metabolism**
**D. EGFR tyrosine kinase inhibitor resistance**
**E. ErbB signaling pathway**
**F. Glutathione metabolism**
**G. mTOR signaling pathway**
**H. PI3K-Akt signaling pathway**
**I. Pyruvate metabolism**

**Supplemental Figure 6. <u>Joint Pathway Analysis in MDA-MB-134 Parent vs FulvR or Parent vs TamR</u>**. All curated pathways identified by JPA with have a p <0.019, false discovery rate (FDR) < 0.055, and a composite score > 1.0.

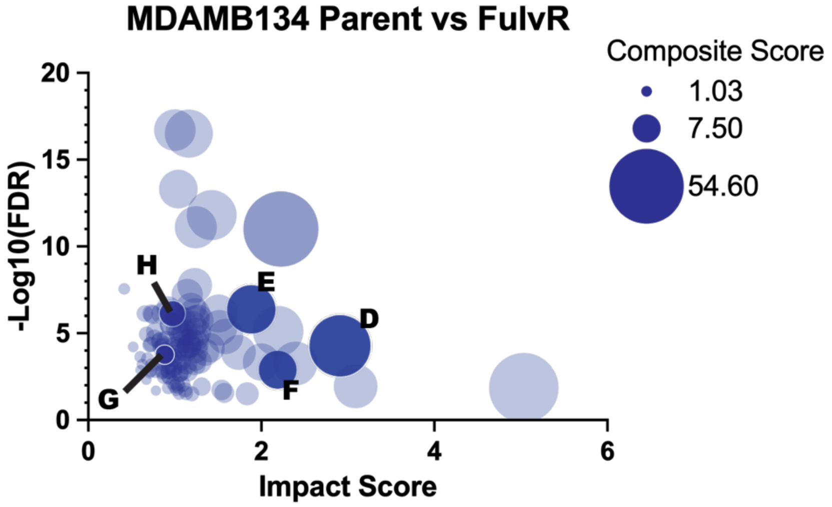

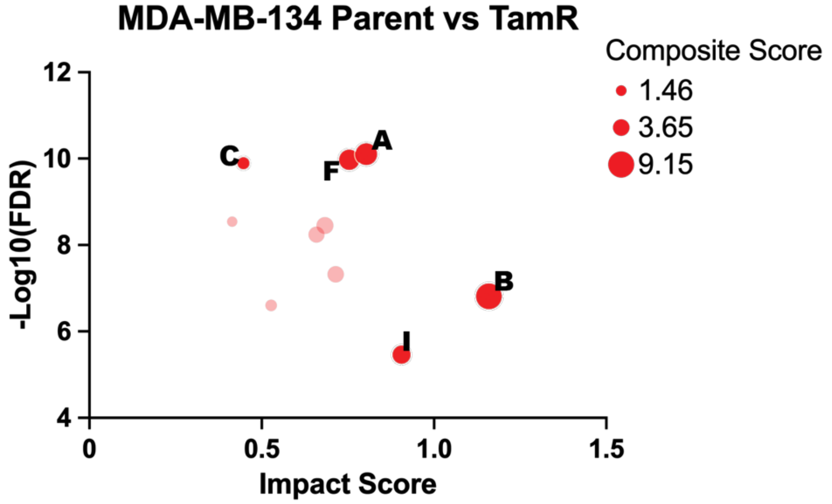

**Key**

**A. Alanine, aspartate and glutamate metabolism**
**B. Citrate cycle (TCA cycle)**
**C. Cysteine and methionine metabolism**
**D. EGFR tyrosine kinase inhibitor resistance**
**E. ErbB signaling pathway**
**F. Glutathione metabolism**
**G. mTOR signaling pathway**
**H. PI3K-Akt signaling pathway**
**I. Pyruvate metabolism**

**Supplemental Figure 7. <u>DMF reverses endocrine resistance in UCD12 parent and TamR cells.</u>** Dimethyl fumarate (DMF) reverses tamoxifen resistance alone and in combination with tamoxifen.

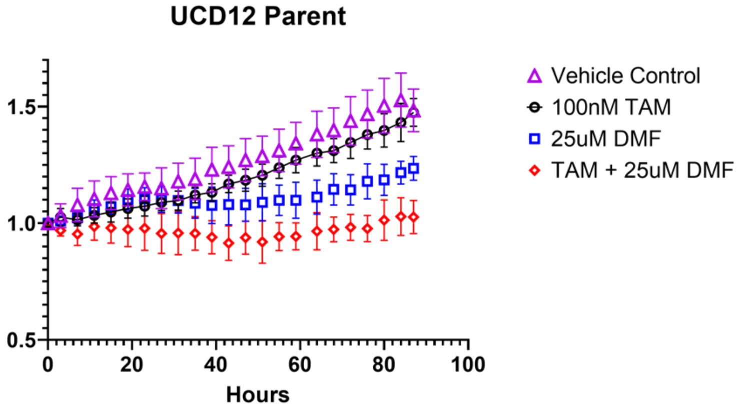

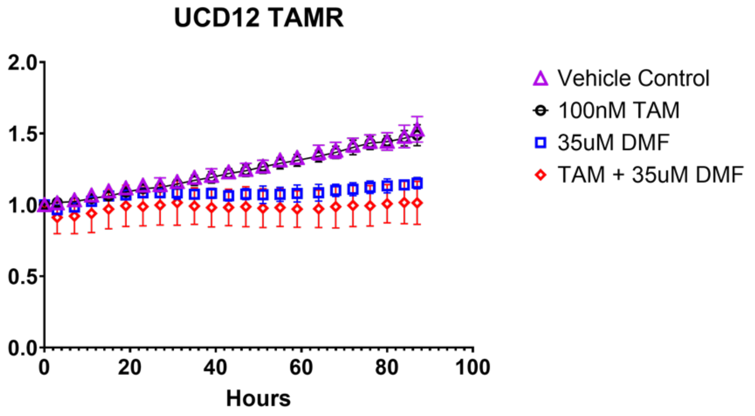

**Supplemental Figure 8. <u>The mevalonate pathway has increased enzymatic expression in the early half of the pathway after DMF treatment in FulvR cells.</u>** Box and whiskers plots of enzymes (proteins) in the mevalonate pathway. Statistical analysis is 1-way ANOVA and significance is $p < 0.05$.

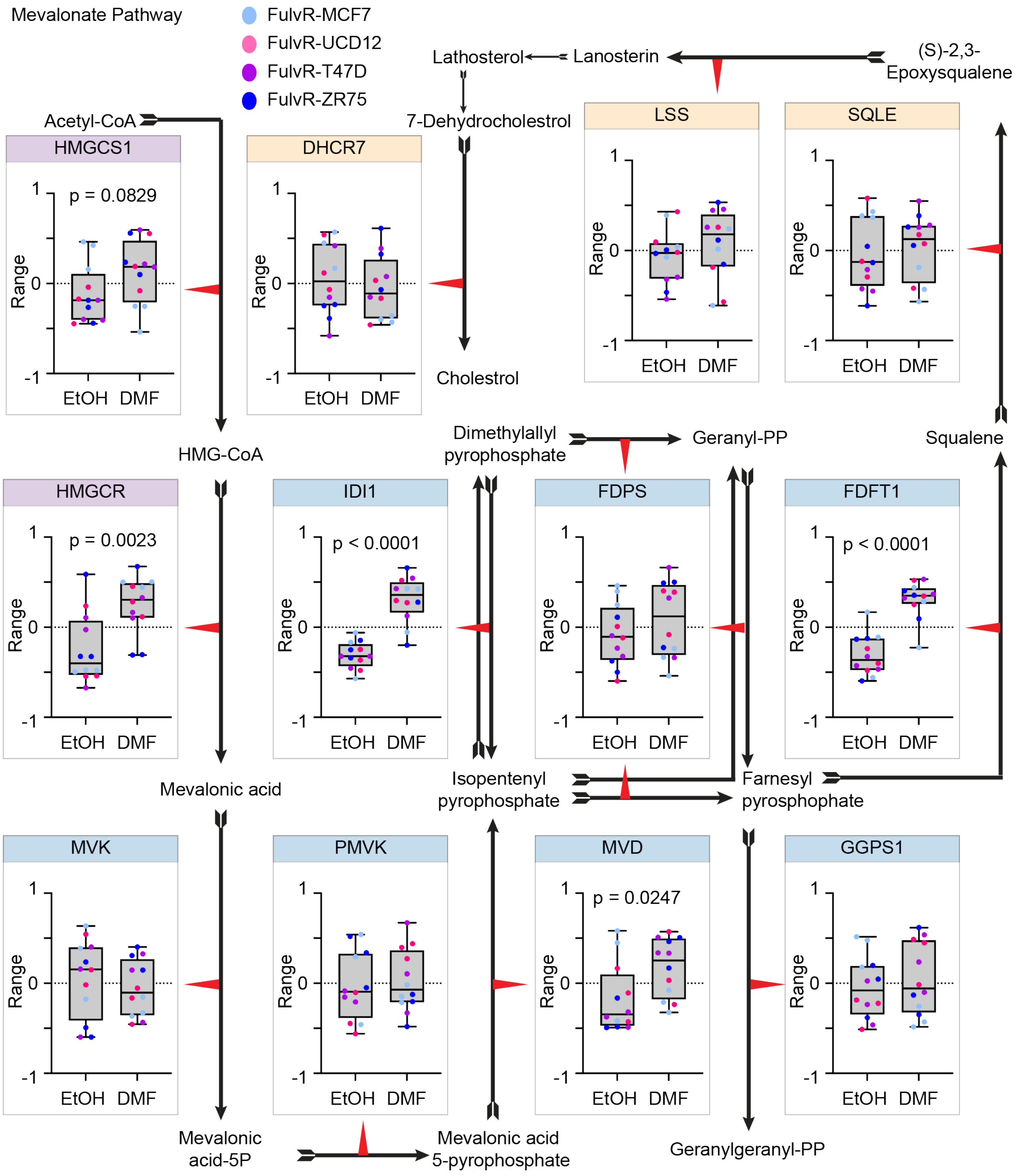

**Supplemental Figure 9.** **The mevalonate pathway has increased enzymatic expression in the cholesterol synthesis half of the pathway after DMF treatment in TamR cells.** Box and whiskers plots of enzymes (proteins) in the mevalonate pathway. Statistical analyses are unpaired t-test with significance is $p < 0.05$.

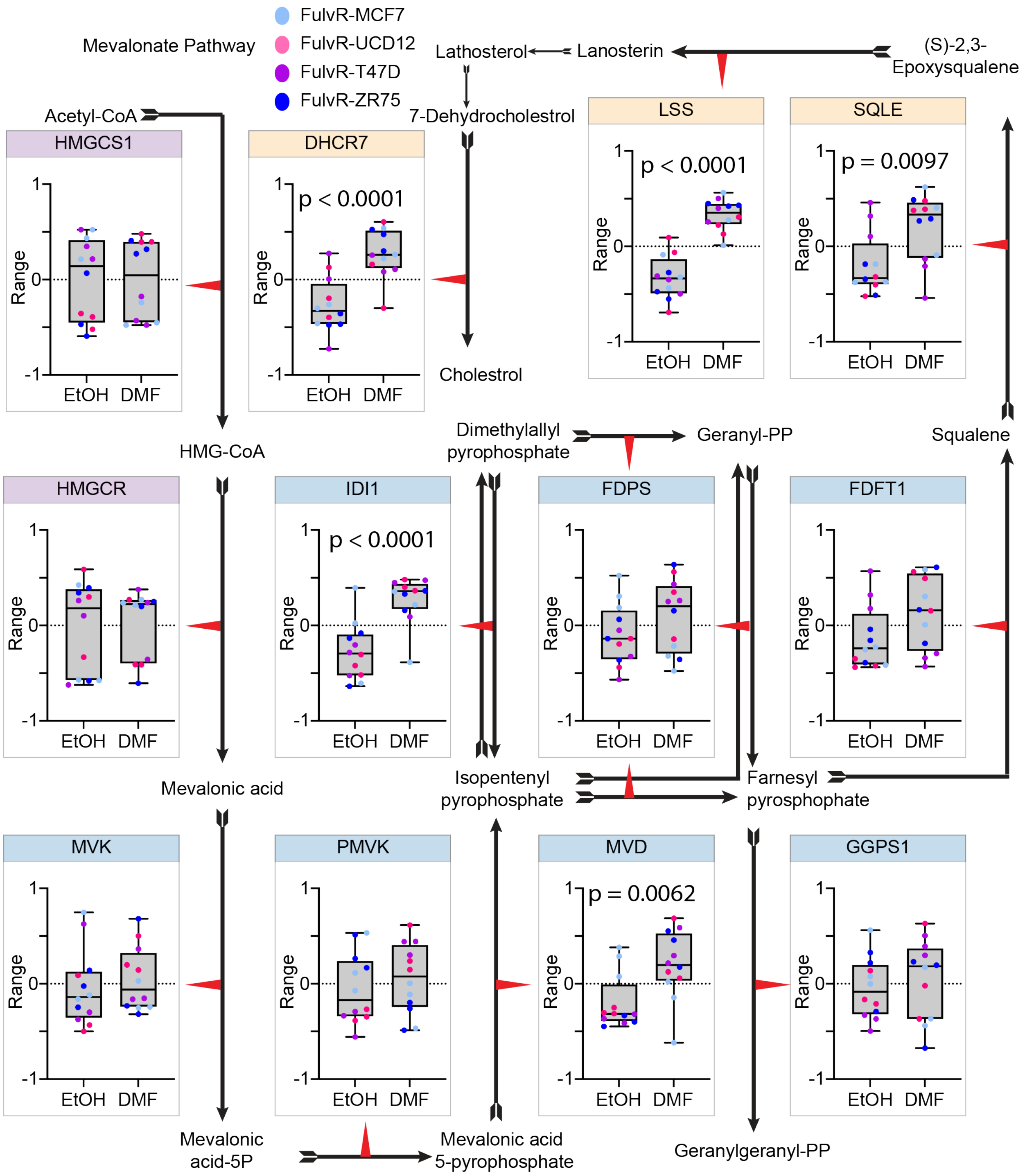

**Supplemental Figure 10. <u>TP53 pathway analysis and protein expression in FulvR cells and TamR cells.</u>** Transcriptional regulation by TP53 is the top pathway identified in RNAseq data of FulvR cells. TP53 protein expression is decreased in FulvR DMF treated cells compared to EtOH treated cells, but there is no difference in expression in TamR cells.

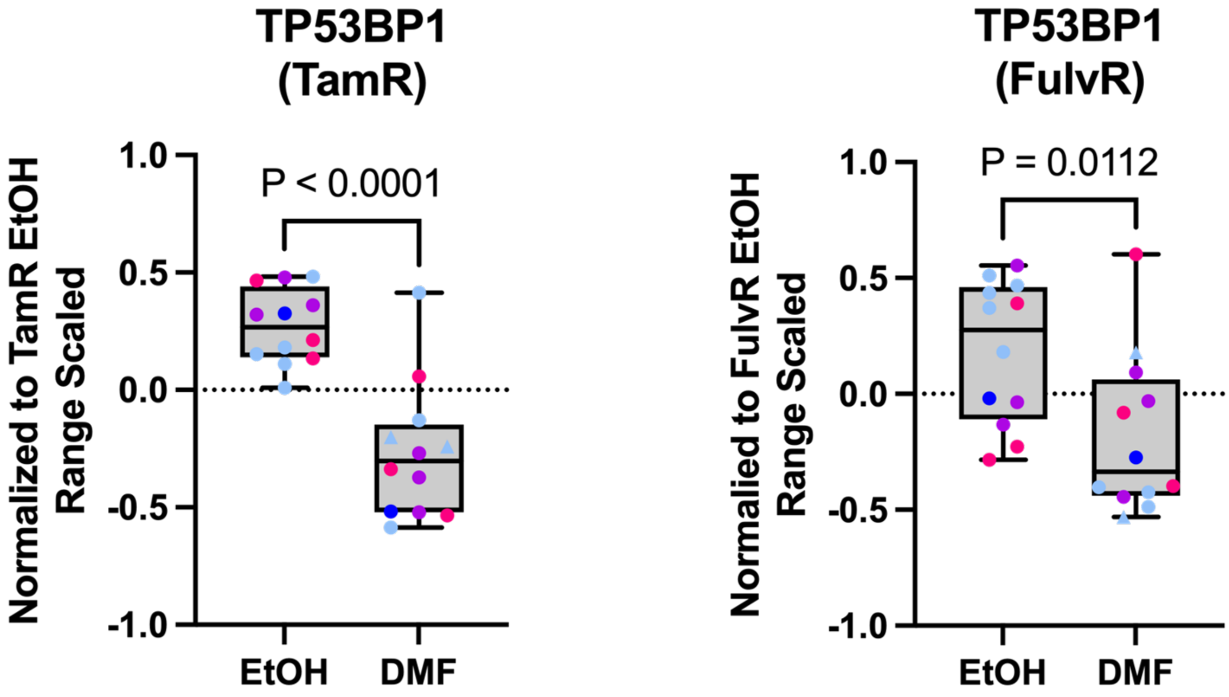

**Supplemental Figure 11.** **Homer motif discovery demonstrates that increased similiarity between Parent EtOH treated and TamR-DMF treated cells.** Venn diagrams of the number of Homer motifs identified in **(A)** Parent EtOH vs TamR EtOH or Parent EtOH vs TamR DMF, and **(B)** chrommHMM-18 marks for the clusters generated by k-means clustering in FulvR and parental cells treated with vehicle or DMF.

**A**

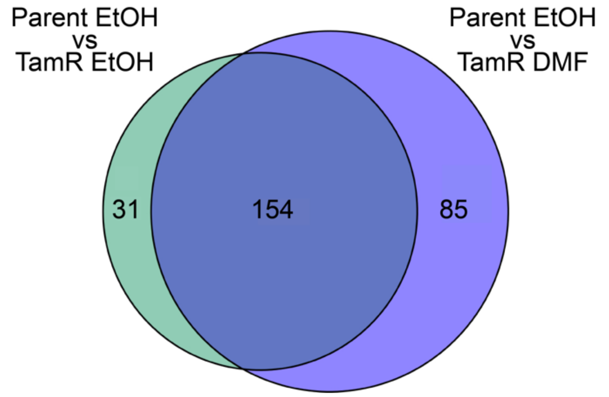

**B**

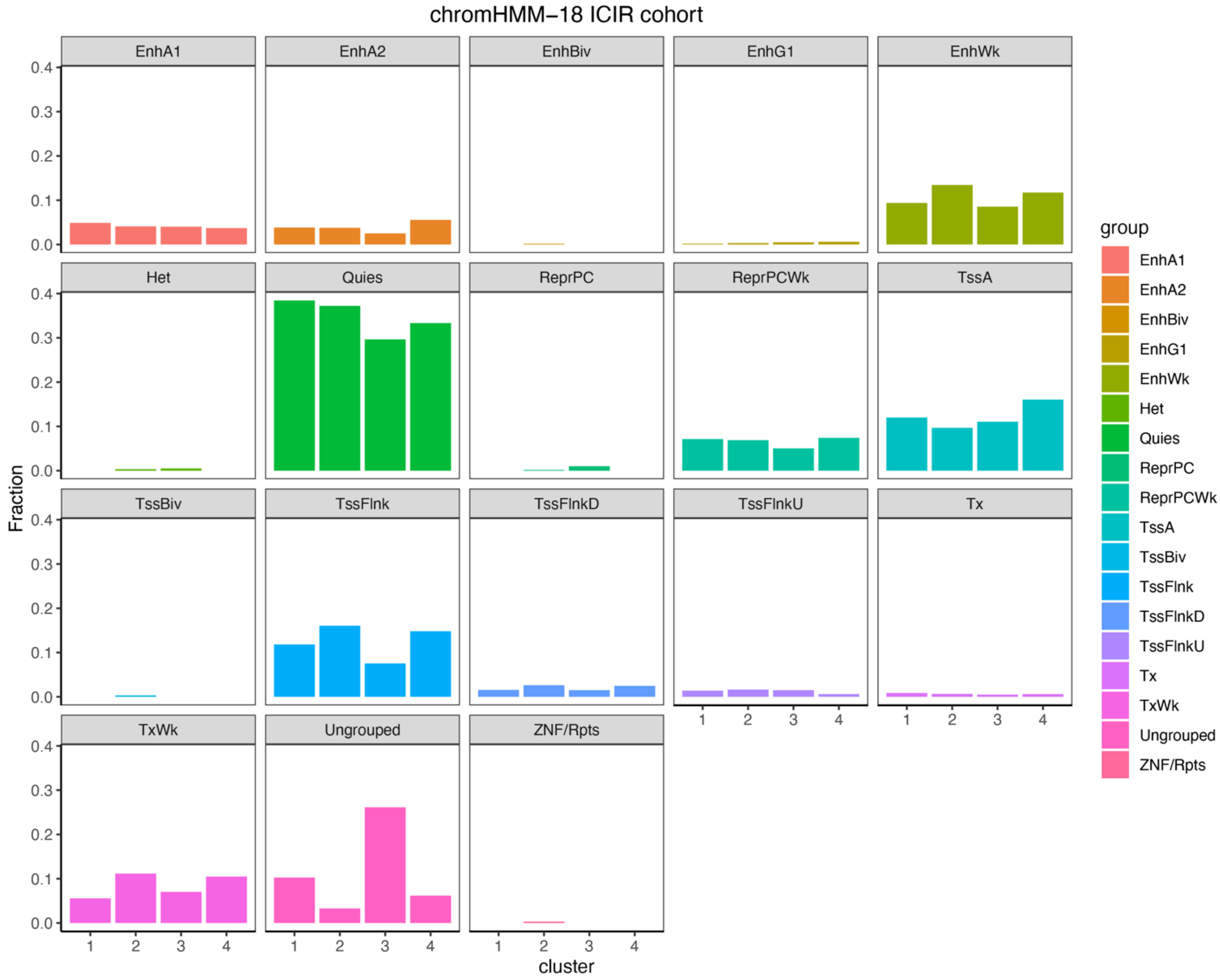

**Supplemental Figure 12. <u>Homer motif discovery demonstrates that increased similiarity between Parent EtOH treated and FulvR-DMF treated cells.</u>** Venn diagrams of the number of Homer motifs identified in **(A)** Parent EtOH vs TamR EtOH or Parent EtOH vs TamR DMF, and **(B)** chrommHMM-18 marks for the clusters generated by k-means clustering in FulvR and parental cells treated with vehicle or DMF.

**Supplemental Figure 12**

**A**

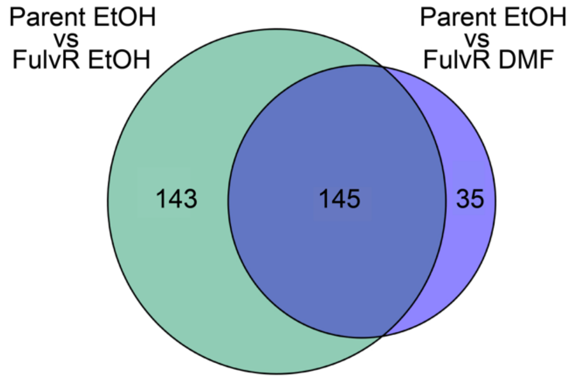

**B**

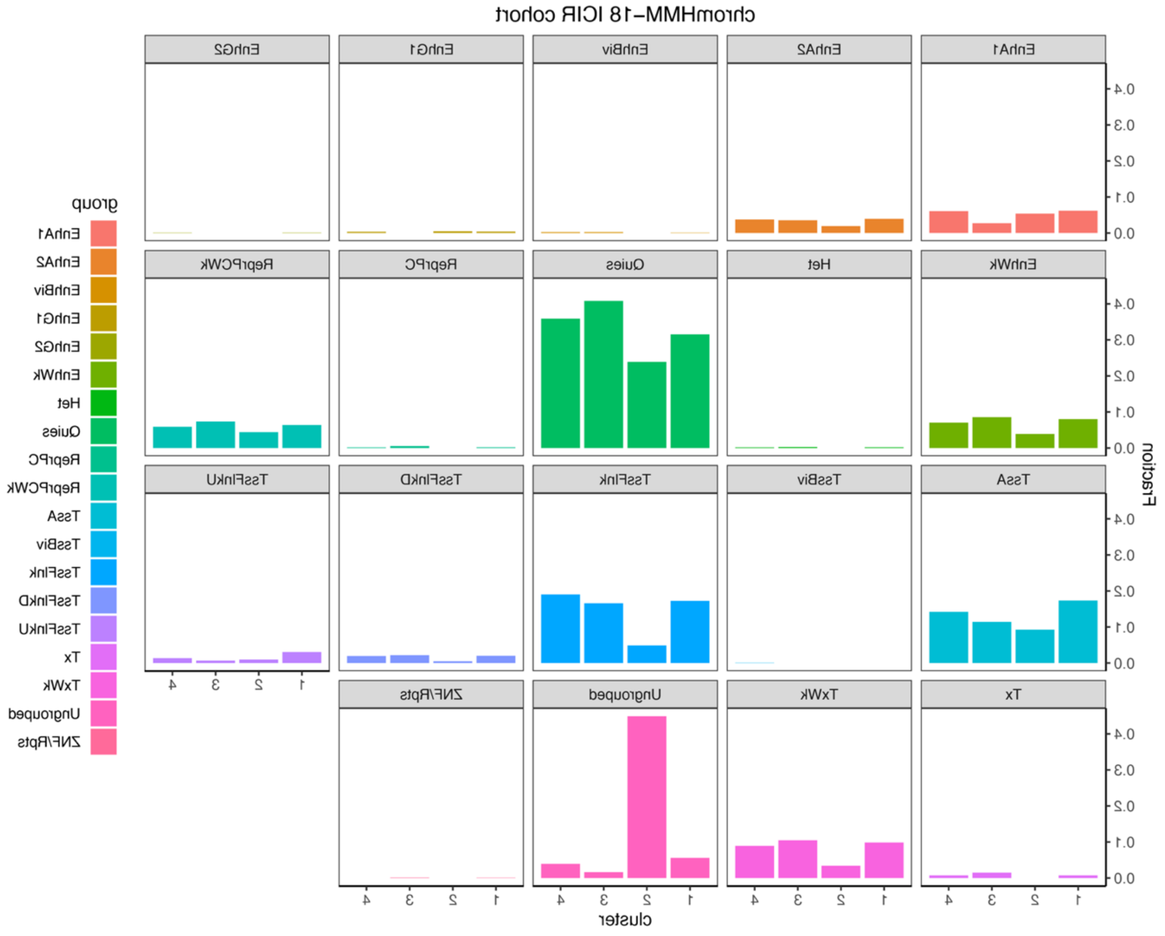

References

1. Finlay-Schultz J, Jacobsen BM, Riley D, Paul KV, Turner S, Ferreira-Gonzalez A, Harrell JC, Kabos P, Sartorius CA. New generation breast cancer cell lines developed from patient-derived xenografts. Breast Cancer Res. 2020;22(1):68. Epub 20200623. doi: 10.1186/s13058-020-01300-y. PubMed PMID: 32576280; PMCID: PMC7310532.
2. Martin M. Cutadapt removes adapter sequences from high-throughput sequencing reads. EMBnetjournal. 2011;17(1):10. doi: 10.14806/ej.17.1.200.
3. Dobin A, Davis CA, Schlesinger F, Drenkow J, Zaleski C, Jha S, Batut P, Chaisson M, Gingeras TR. STAR: ultrafast universal RNA-seq aligner. Bioinformatics. 2013;29(1):15-21. Epub 20121025. doi: 10.1093/bioinformatics/bts635. PubMed PMID: 23104886; PMCID: PMC3530905.
4. Love MI, Huber W, Anders S. Moderated estimation of fold change and dispersion for RNA-seq data with DESeq2. Genome Biology. 2014;15(12). doi: 10.1186/s13059-014-0550-8.
5. Langmead B, Salzberg SL. Fast gapped-read alignment with Bowtie 2. Nature Methods. 2012;9(4):357-9. doi: 10.1038/nmeth.1923.
6. Li H, Handsaker B, Wysoker A, Fennell T, Ruan J, Homer N, Marth G, Abecasis G, Durbin R. The Sequence Alignment/Map format and SAMtools. Bioinformatics. 2009;25(16):2078-9. Epub 20090608. doi: 10.1093/bioinformatics/btp352. PubMed PMID: 19505943; PMCID: PMC2723002.
7. Li H. A statistical framework for SNP calling, mutation discovery, association mapping and population genetical parameter estimation from sequencing data. Bioinformatics. 2011;27(21):2987-93. Epub 20110908. doi: 10.1093/bioinformatics/btr509. PubMed PMID: 21903627; PMCID: PMC3198575.
8. Lloyd S. Least squares quantization in PCM. IEEE Transactions on Information Theory. 1982;28(2):129-37. doi: 10.1109/tit.1982.1056489.
9. Bailey TL, Johnson J, Grant CE, Noble WS. The MEME Suite. Nucleic Acids Research. 2015;43(W1):W39-W49. doi: 10.1093/nar/gkv416.
10. Grant CE, Bailey TL, Noble WS. FIMO: scanning for occurrences of a given motif. Bioinformatics. 2011;27(7):1017-8. Epub 20110216. doi: 10.1093/bioinformatics/btr064. PubMed PMID: 21330290; PMCID: PMC3065696.
11. Xia J, Psychogios N, Young N, Wishart DS. MetaboAnalyst: a web server for metabolomic data analysis and interpretation. Nucleic Acids Res. 2009;37(Web Server issue):W652-60. Epub 20090508. doi: 10.1093/nar/gkp356. PubMed PMID: 19429898; PMCID: PMC2703878.
